# Sequence-dependent scale for translocon-mediated insertion of interfacial helices in membranes

**DOI:** 10.1101/2024.10.23.619793

**Authors:** Brayan Grau, Rian Kormos, Manuel Bañó-Polo, Kehan Chen, María J. García-Murria, Fatlum Hajredini, Manuel M. Sánchez del Pino, Hyunil Jo, Luis Martínez-Gil, Gunnar von Heijne, William F. DeGrado, Ismael Mingarro

## Abstract

Biological membranes consist of a lipid bilayer studded with integral and peripheral membrane proteins. Most α-helical membrane proteins require protein-conducting insertases known as translocons to assist in their membrane insertion and folding. While the sequence-dependent propensities for a helix to either translocate through the translocon or insert into the membrane have been codified into numerical hydrophobicity scales, the corresponding propensity to partition into the membrane interface remains unraveled. By engineering diagnostic glycosylation sites around test peptide sequences inserted into a host protein, we devised a system that can differentiate between water-soluble, surface-bound, and transmembrane (TM) states of the sequence based on its glycosylation pattern. Using this system, we determined the sequence-dependent propensities for transfer from the translocon to a TM, interfacial or extramembrane space. UMAP analysis of a large collection of TM and water-soluble helices provide useful embeddings for analysis of these propensities and aid in understanding the physical properties and functions of antimicrobial, lytic, and fusogenic peptides.

## Introduction

While the sequence characteristics required for translocon-mediated insertion and stabilization of transmembrane (TM) helices in membrane proteins have been extensively studied for decades ^1^, there has been a paucity of corresponding data examining the insertion of sequence elements such as amphipathic α-helices into the membrane interface. The dynamic equilibrium between bulk water, the membrane surface and TM states represents a delicate balance that is essential for the function of peptides such as antimicrobial peptides (AMPs) and lytic peptides. Moreover, proteins with helical fusogenic sequences similarly partition between water-soluble, membrane-associated, and TM fusion pore-forming states ^2,3^. Other surface-interacting peptides are known to stabilize membrane curvature ^4^. Thus, elucidating the features that dictate membrane surface versus TM associations is essential to a wide swath of natural proteins ^5,6^.

Biological membranes can be divided into two regions based on physicochemical properties: the highly hydrophobic core formed mainly by the lipid acyl chains and the interfaces on both sides of this central region that contain the polar head groups^7^. The hydrophobic effect is the primary driving force of membrane partitioning to the former, but much less is known about the energetics of interface partitioning. The combined thickness of the interfaces is similar to the thickness of the hydrophobic core (∼30Å), and this region is able to accommodate unfolded and folded polypeptide chain ^8^. The interfacial region, occupied by the lipid headgroups and the associated hydration layer, is highly physically anisotropic. Because the interfaces are rich in groups with different chemical properties, a polypeptide chain in this region faces an environment enriched in possibilities to establish a variety of noncovalent interactions.

Translocons have evolved to recognize and insert hydrophobic TM segments into the bilayer in a manner that optimizes side chain interactions with both the hydrophobic core and interfacial regions ^6,9^. Structural data have provided detailed insights concerning the insertion into the hydrophobic sector of the bilayer, showing that TM segments leave the translocon through a lateral gate in the channel wall that opens laterally toward the bilayer ^10–13^. The precise mechanistic details behind the recognition of interfacial sequences, however, remain obscure.

To achieve a quantitative description of membrane protein insertion and folding, it becomes necessary to unravel the molecular processes by which a polypeptide segment exiting the translocon adopts a TM orientation (spanning the membrane) versus sliding toward the membrane surface in an interfacial disposition ^6,14^. Once a detailed description of TM segment recognition by the translocon has been established ^15,16^, a fundamental requirement for a quantitative characterization of protein sliding and folding in membrane interfaces is a suitable interfacial hydrophobicity scale. In this study, we present such a scale for the twenty natural amino acids derived from quantitative measurements of the translocon-mediated protein integration pathway into the ER membrane.

To develop this scale, we challenged the translocon both *in vitro* and in cellular membranes with a set of designed polypeptide sequences. Our designs are based on a peptide sequence derived from bacteriorhodopsin helix C (bRc) that out of the protein context does not insert into a lipid bilayer as a membrane-spanning helix at physiological pH but rather adopts a surface configuration with a high helical content in the presence of DOPC liposomes ^17^. We then substituted different amino acids into this peptide sequence to modulate the energetics of partitioning into the TM versus surface state. Through rational design of glycosylation sites, we devised a system that can differentiate between water-soluble, surface-bound and TM states of the peptide sequence based on its glycosylation pattern. A quantitative analysis of the mole fraction of the protein in each state allows one to compute an apparent free energy of transfer (ΔG_app_) from water to both the surface-absorbed and TM states for each of the twenty natural amino acids.

We compare the data from our novel ΔG_app_ scales with that obtained from 1) previous studies of translocon-mediated TM insertion ^15,16^; 2) biophysical measurements ^18,19^ of partitioning into organic solvents and the surface of phospholipid bilayers; and 3) previous ^20^ statistical analyses of the depth-dependent distribution of amino acids in membrane proteins of known structures. Our biologically and statistically derived scales show reasonable agreement with biophysical scales, with some interesting exceptions associated with the tendency of basic and aromatic residues to associate with the interfacial region of membrane bilayers. These sequence-specific propensities for the interface, even among residues that are charged or have expansive hydrophobic surface area, appear to enable more precise targeting of helices to the interface than could be achieved by merely selecting sequences with intermediate hydrophobicity. The derived scale is then shown to be useful for analysis of membrane-associated helical proteins.

## Results

### Development of an assay to assess TM, interfacial, and water-soluble orientations in a single experiment

Previously, Hessa and coworkers developed a “biological” hydrophobicity scale based on an *in vitro* assay ^15,16^ for quantifying the efficiency of translocon-mediated membrane integration of TM helices into dog pancreas rough microsomes (RMs). In this method, a test sequence is engineered into the luminal P2 domain of the integral membrane protein leader peptidase (LepB), where it is flanked by two acceptor sites for N-linked glycosylation (Fig. 1a). The degree of membrane integration of the test sequence is quantified from SDS–PAGE gels by measuring the fraction of singly versus doubly glycosylated LepB molecules. If the test sequence is inserted into the membrane adopting a TM orientation, then only one site (designated the G1 site) is glycosylated, while double glycosylation at G1 and G2 (or G2’) is observed when the test sequence fails to insert into the membrane and instead is translocated to the lumen. Here, we wished to additionally probe the sequence requirements for the test sequence to associate tightly with the luminal surface of the membrane. We hypothesized that a G2 site proximal to the test sequence would be protected from glycosylation if the test sequence associated tightly to the membrane ^21,22^. As a test sequence we used a bacteriorhodopsin helix c derived (bRc) interfacial peptide, developed by Engelman et al.^17^, which is helical in the presence of POPC liposomes (Supplementary Fig. S1) and, depending on conditions, capable of adopting a TM or interfacial conformation maintaining more amphiphilic character than the wild type bRc sequence.

**Figure 1.**
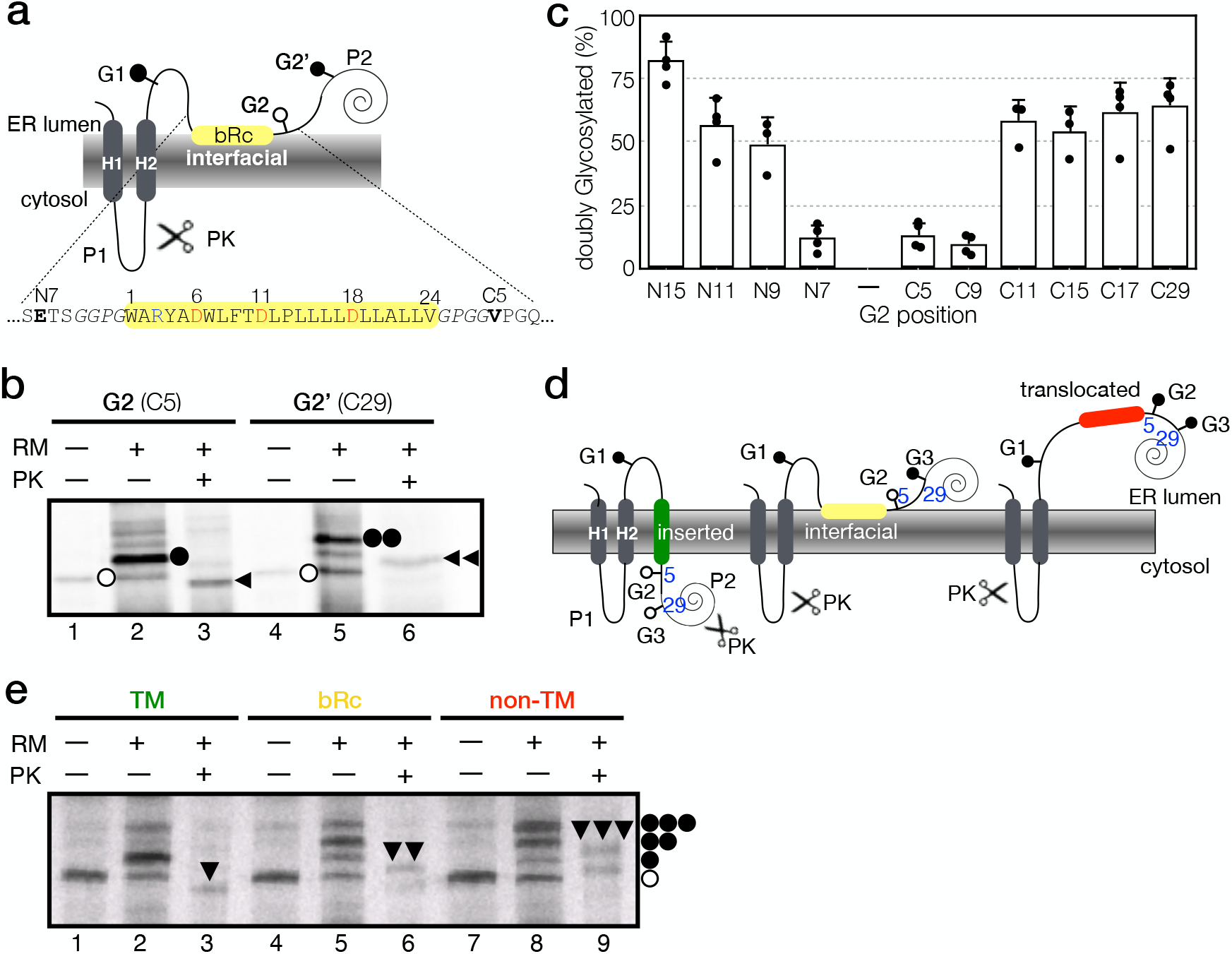
Interfacial segment disposition in the microsomal membrane via a glycosylation-based assay. **a**. Schematic representation of the modified glycosylation-based assay for discriminating between TM and aqueous peptide dispositions, introduced by Hessa and coworkers ^15,16^. Wild type Lep has two N-terminal TM segments (H1 and H2) and a large luminal domain (P2) in which interfacial segment was inserted (yellow highlighted). Glycosylation sites are inserted at positions both N-terminal and C-terminal to the test peptide and serve as indicators of the topology in which the full translation product has inserted into the membrane. G1 is always placed in positions 96-98 of the wild type LepB and C-terminal glycosylation acceptor sites were placed next to the insulating GPGG sequence (G2) or 29 residues C-terminal from the insulating sequence (G2’), displaying a glycosylation pattern regarding the final disposition of the tested sequence. **b**. Plasmids encoding the Lep/bRc constructs were transcribed and translated *in vitro* in the presence (+) and absence (-) of dog pancreas rough microsomes (RM) and proteinase K (PK). Tested sequence bRc is highlighted in a yellow box. Bands of non-glycosylated protein are indicated by a white dot; singly and doubly glycosylated proteins are indicated by one or two black dots, respectively. The arrowhead identifies undigested protein after PK treatment. One and two arrowheads indicate singly and doubly glycosylated protected fragments, respectively. **c**. Determination of the minimal glycosylation distance for interfacial sequences. Glycosylation site G2 were moved across different positions in both N-terminal (N15, N11, N9, and N7) and C-terminal (C5, C9, C11, C15, C17, and C29) of the tested sequence bRc. – represent the absence of glycosylation site G2. Error bars show the standard deviation of 3 or more independent experiments. Black and white rectangles represent when a tested sequence is or is not mainly represented by doubly glycosylated molecules, respectively. **d**. Schematic representation of the LepG3 design where all three glycosylation acceptor sites (G1, G2, and G3) are placed in the same construct, displaying a glycosylation pattern reflecting the final disposition of the assayed sequence (inserted, surface, or translocated). **e**. *In vitro* translation in the absence (—) or presence (+) of rough microsomes (RM) of LepG3 constructs harboring the first TM segment for *Turnip Crinkle Virus* movement protein (TM)^26^, bRc derived interfacial sequence, and a translocated pseudo-randomized sequence from Ref. ^46^. Bands of non-glycosylated proteins are indicated by a white dot; single, double, and triple glycosylated proteins are indicated by one, two, or three black dots, respectively. The protected double or triple glycosylated H2/G1/bRc/G2/G3/P2 and H2/G1/translocated/G2/G3/P2 fragments from proteinase K assay (PK) are indicated by two or three arrowheads, respectively.

We thus began by testing the bRc sequence flanked by GGPG and GPGG tetrapeptides to insulate the interfacial segment from the surrounding sequence ^15^, placing the G2 site at two different positions. We first placed the G2 site only 5 residues after the test sequence such that it would not be accessible to the luminal oligosaccharyl transferase (OST) active site ^21,23^ if the test sequence partitioned into the interfacial region of the membrane. In this construct, with bRc as the test sequence, the G2 site was protected from glycosylation, while the G1 site remained efficiently glycosylated (Fig. 1b, lane 2). Proteinase K digestion revealed a luminal disposition of the test sequence, with the LepB P2 domain protected from degradation (Fig. 1b, lane 3). Moving the second glycosylation site further away from the tested sequence (G2’) yielded mainly doubly glycosylated molecules resistant to proteinase K treatment (Fig. 1b, lanes 5 and 6), confirming the luminal location of the P2 domain. These results were confirmed using a second peptide, melittin, known to form an interfacial helix ^24,25^ as the test sequence (Supplementary Fig. S2), which was also mostly protected from glycosylation at G2, accessible for glycosylation at G2’, and with the P2 loop protected from proteinase K degradation.

To further validate the assay, the “minimal glycosylation distance” for interfacial sequences was measured by placing the G2 site at different positions upstream and downstream the bRc sequence (Supplementary Fig. S3). We found that efficient glycosylation was observed when the Asn residue in the G2 site was placed at least 11 residues downstream (Figs. 1c and S3), and at least 9 residues upstream of the bRc sequence (Figs. 1c and S3).

In all subsequent experiments, we kept the G2 site 5 residues after the test sequence and introduced a third site, G3 (Fig. 1d), 29 residues away from the test sequence (the farthest tested position in the minimal glycosylation distance screening) to be able to determine whether the C-terminus of the construct was luminal (as in a surface-absorbed state) or cytoplasmic (as in the TM state). With these three sites (G1, G2, and G3) simultaneously present (LepG3), the glycosylation state can be used to determine whether the test sequence in the translation product is TM, interfacial, or soluble (Fig. 1d). In the TM state, the G2 and G3 sites locate to the cytosol, and hence only the G1 site is glycosylated, leading to a singly glycosylated band on an SDS-PAGE electrophoresis gel. If the test sequence instead associates tightly with the luminal membrane surface, a doubly glycosylated product is observed due to modifications at G1 and G3. Finally, a triply glycosylated product is the predominant product when the test sequence is fully translocated into the luminal space. Indeed, the expected single-glycosylation pattern was observed when the test sequence was substituted with a known TM sequence ^26^ (Fig. 1e, TM, lane 2), and the triply glycosylated pattern predominates when a polar C-terminal peptide sequence from LepB served as the test sequence (Fig. 1e, non-TM, lane 8). Moreover, as expected, the bRc sequence gave a mixture of singly, doubly and triply glycosylated products, with doubly glycosylated molecules being the most abundant (Fig. 1e, bRc, lane 5). Protection of the P2 domain from proteinase K digestion confirmed this membrane disposition (Fig. 1e, lane 6).

We also challenged our assay with a set of 19-residue test sequences composed entirely of Ala and Leu ^15^, with compositions ranging from two to five Leu residues. This family of sequences showed a clear transition from triple to single glycosylation as the number of Leu residues was increased (Supplementary Fig. S4), showing that the assay using model Ala and Leu hydrophobic sequences can resolve the state preferences of sequences that partition into a mixture of soluble and TM states, but not the interfacial state.

### bRc-derived sequence as a vehicle for amino acid contribution

In order to develop a strategy to measure the effect of amino acid substitutions on membrane integration, we next modified the test sequence from the bRc peptide to favor either TM or fully exposed (luminal) conformations. The interfacial location of bRc is believed to result from three Asp residues, D6, D11 and D18 (Fig. 1a) and single substitutions within this peptide changed its disposition in previous studies with synthetic peptides ^17^ and *in vitro* transcription/translation experiments ^27^. In agreement with the earlier data, we found that the D11L and D18L variants, in which a single charged Asp is changed to an apolar Leu, becomes fully transmembrane (i.e., the singly glycosylated band predominates and no proteinase K protection is observed, Supplementary Fig. S5, lanes 2, 3, 11 and 12, in contrast to the WT sequence in Fig. 1b, lanes 2 and 3). Conversely, replacing an existing Leu with an Asp (variants L12D and L17D) shifts the glycosylation pattern to the doubly glycosylated, translocated state with the P2 domain protected from proteinase K degradation (Supplementary Fig. S5, lanes 5, 6, 14 and 15, respectively). Interestingly, when both Asp and Leu residues in positions 11/12 or 17/18 were replaced by Trp residues (D11W/L12W or L17W/D18W), which strongly prefer the membrane interface ^18^, we observed singly-glycosylated forms and PK treatment protection (Supplementary Fig. S5b, lanes 7-9 and 16-18, respectively), supporting a primarily interfacial disposition. Those results confirm that the bRc-derived sequence is able to provide a suitable vehicle to study the contribution of any single amino acid along its sequence, being its final membrane disposition successfully switched with punctual mutations. Ultimately, we chose the more centered 11 and 12 positions over the more peripheral 17 and 18 positions, because the latter bias the initiation of helix formation ^28^.

To prove the suitability in our LepG3 assay of bRc-derived sequence as a backbone to study the contribution of single amino acids, we substituted Asp at position 11 with Leu and Ala (Supplementary Fig. S6, constructs #1 and #2), which increased the fraction in the transmembrane-inserted state. We next increased the number of Asp residues in the peptide by substituting the hydrophobic residues at positions 12-17 with increasing numbers of Asp residues (Supplementary Fig. S6, constructs #4 to #7). The consecutive aspartate substitutions shift the banding pattern towards the fully exposed, triply glycosylated state, reaching a maximum after only two Asp substitutions are introduced, which creates a triplet of Asp residues considering the presence of the original D11. This result again shows the sensitivity of the system to small changes.

We next compared the effect of replacing the original residues at the center of the bRc-derived sequence with triplets of amino acids. Triplet substitutions were chosen to maximize the effects and because they displayed a clearer glycosylation pattern than pair-substitutions (Supplementary Fig. S7). For the hydrophobic leucine triplets, the singly-glycosylated form was predominant (Supplementary Fig. S7c, lane 2), indicative of a TM disposition, whereas the tryptophan triplet mainly produced doubly-glycosylated molecules (Fig. S7c, lane 5), indicating a surface location. Finally, the construct harboring three Asp residues yielded triply-glycosylated bands (Fig. S7c, lane 8) as expected for the fully exposed, luminal orientation of this sequence. We therefore chose this system for a systematic study of the effects of substitutions on membrane association and insertion.

### Biological interfacial and transmembrane scales

We used triplet substitutions in the bRc-derived sequence to quantify the mole fractions of the three glycosylation states, from which we computed the apparent free energies of transfer of each of the 20 naturally occurring amino acid sidechains from water to the membrane interface (ΔG_app_^Wat→Int^) or to the hydrophobic core of the membrane (ΔG_app_^Wat→TM^). First, we investigated the contribution of each position within the triplet, results obtained when 1-3 Gly residues are incorporated on an Ala triplet showed an overall linear relationship between the number of Gly residues and ΔG_app_, a simple outcome consistent with energy-additivity (Supplementary Fig. S8). As described in Methods, we evaluated a total of 112 measurements on 27 variants using a rabbit reticulocyte-derived translation system to obtain the biologically derived *in vitro* scale. As expected, the ΔG_app_^Wat→TM^ values obtained from 3-state system (Fig. 2a, Supplementary Table 1) correlate well with the depth-dependent insertion scale of Hessa et al. ^16^. The ΔG_app_^Wat→TM^ scale correlates best with the Hessa (HWvH) values for transfer of an amino acid to locations near the center of the bilayer (R^2^ = 0.78 to 0.79 within the range spanning ± 4.5 Å of the center, Fig. 2b and 2d). By contrast, the ΔG_app_^Wat→Int^ values correlate best with the corresponding values for transfer of a sidechain from water to a depth more consistent with the boundary between the bilayer core and the interface regions (R^2^ = 0.79-0.81 within the pair of ranges spanning 7.5 ± 1.5 Å and -7.5 ± 1.5 Å from the center, Fig. 2c and 2d). These findings are in good agreement with our expectation that ΔG_app_^Wat→TM^ values should correlate with the center of the bilayer, while ΔG_app_^Wat→Int^ values should reflect transfer to a more interfacial location in the bilayer.

**Figure 2.**
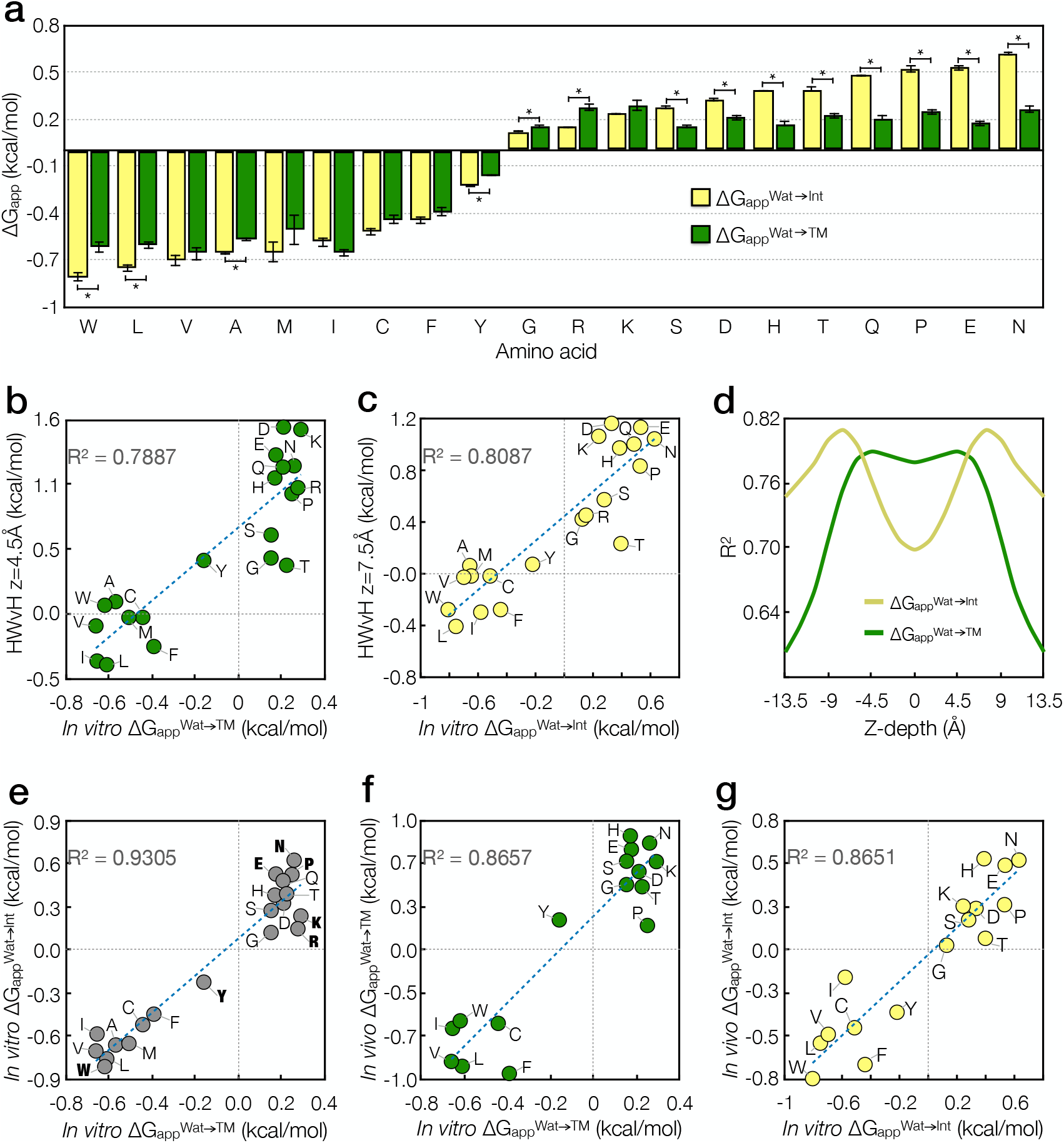
Biological and biophysical ΔG_app_ scales. **a**. ΔG_app_ of transferring for each amino acid type from the aqueous translocon channel to the membrane interface (ΔG_app_^Wat→Int^, yellow bars) and to the hydrophobic core of the membrane (ΔG_app_^Wat→TM^, green bars). Amino acids for which the difference between ΔG_app_^Wat→Int^ and ΔG_app_^Wat→TM^ was statistically significant are marked with asterisks. **b**. Correlation between ΔG_app_^Wat→TM^ and the position-dependent scale of Hessa et al. ^15,16^ at a position ±3 residues, or equivalently 4.5 Å, from the center of the bilayer. **c**. Correlation between ΔG_app_^Wat→Int^ and the position-dependent scale of Hessa et al. ^9,10^ at a position ±5 residues, or equivalently 7.5 Å, from the center of the bilayer. **d**. Coefficient of determination (R^2^) values for linear least squares trendlines between all position-dependent scales of Hessa et al. ^16^ and ΔG_app_^Wat→Int^ (yellow line) or (ΔG_app_^Wat→TM^ (green line). **e**. Correlation between ΔG_app_^Wat→TM^ and ΔG_app_^Wat→Int^. Amino acids with major dispersion of the trend line are marked in bolt.**f**. Correlation between *in vivo* ΔG_app_^Wat→Int^ scale and *in vitro* ΔG_app_^Wat→Int^ scale. **g**. Correlation between *in vivo* ΔG_app_^Wat→TM^ scale and *in vitro* ΔG_app_^Wat→TM^ scale.

The current study represents the first experimental scale for the effects of substitutions on an interfacially oriented versus a TM inserted helical sequence under identical experimental conditions. Thus, it is of interest to compare the interfacial and transmembrane scales (Supplementary Table 1). A plot of ΔG_app_^Wat→Int^ vs. ΔG_app_^Wat→TM^ (Fig. 2e) shows that the two scales are well correlated (R^2^ = 0.93), as anticipated from the fact that both are measures of the transfer of a sidechain from water to a more apolar environment. Interestingly, the trendline has a slope near unity (1.27), indicating that ΔG_app_^Wat→Int^ is similar in magnitude to ΔG_app_^Wat→TM^.

Despite the high overall correlation between ΔG_app_^Wat→Int^ and ΔG_app_^Wat→TM^, closer comparison of the two scales reveals important information about the particular amino acids (Fig. 2e, bold annotations) that most strongly favor an interfacial versus transmembrane disposition. The positively charged residues Arg and Lys showed the largest deviation from the regression trendline, favoring the interfacial region over a transmembrane orientation. This finding is in good agreement with their positive charge, which is complementary to the anionic membrane lipids. Consistent with this observation, the anionic residues Asp and Glu have less favorable values of ΔG_app_^Wat→Int^ compared to Arg and Lys (Fig. 2a), though only Glu shows a significant deviation from the trendline (Fig. 2e). Tyr and Trp also show significant deviations favoring the surface orientation, as they have an amphiphilic structure with a polar OH or indole NH, respectively, connected to an otherwise apolar aromatic core sidechain. Amphiphilicity would appear to be more important than aromatic character, as Phe does not deviate strongly from the trendline. Finally, helix-breaking residues such as Pro and Asn are destabilizing to a surface association. This finding might be at least partially a result of the coil-to-helix orientation that accompanies the binding of peptides to the membrane interface ^29–31^, which is expected to be reflected in the coupled energetics associated with such binding. By contrast, a helix is the default orientation for sequences as they are transmitted from the translocon to a TM orientation, and even Pro substitutions are easily accommodated in the TM helices of membrane proteins ^22,32^.

### Development of biological scales for ΔG_app_^Wat→TM^ and ΔG_app_^Wat→Int^ based on expression in mammalian cells

While the above studies were conducted using an *in vitro* assay, it was important to ascertain that the results extend to a cellular milieu. To address this question, we tagged appropriated control sequences plus 17 representative variants of bRc and expressed them in HEK293 cells to measure an *in vivo* scale. As shown in Figure S9a, an unambiguous glycosylation pattern arises, with the construct harboring the TM sequence being singly-glycosylated, the one encoding the bRc variant sequence being doubly-glycosylated, and the construct harboring a non-TM sequence being triply-glycosylated, denoting an inserted, interfacial and translocated location, respectively. As before, triplet substitutions were chosen to maximize the effects and because they displayed a clearer glycosylation pattern than pair-substitutions in mammalian cells (Supplementary Fig. S9b and S9c). The *in vitro* and *in vivo* scales are highly correlated (R^2^ = 0.87 for both, Wat→Int and Wat→TM, Fig. 2f and 2g, respectively), and the deviations from linearity were largely within the experimental error seen in the individual *in vitro* measurements (Fig. 2a). These deviations can also arise from the presence in the cellular system of targeting factors and/or insertase components not included in the *in vitro* microsomal membranes. These findings indicate that the energetics measured *in vitro* are quite predictive of those observed in a cellular milieu.

### Comparison of biological interfacial and transmembrane scales with scales derived from structural informatics

Biologically derived hydrophobicity scales ^16^ have previously been shown to correlate well with knowledge-based scales derived from the frequency of occurrence of the 20 amino acid sidechains in membrane proteins, although the absolute range of the free energies obtained are reduced for knowledge-based versus biologically derived scales. It has been suggested that this attenuation arises from a lack of consideration of interior versus exposed positions during derivation of statistical potentials ^33^. The increase in the number of structures of membrane proteins in recent years has now allowed us to calculate the propensities of each amino acid as a function of both lipid-exposure and depth in the lipid bilayer. A total of 2,229 membrane proteins (1,159,085 residues) were analyzed and their positions within the bilayer (designated z-positions) were taken from the assignments in the OPM database ^34^. The propensities for each residue type varied considerably based on both their membrane depth and surface-accessibility. We used these propensity scales to calculate the apparent free energy required to move an amino acid from an exposed water-soluble state to varying depths using the reverse Boltzmann approximation as described previously ^33,35^. The center of the bilayer was designated as z = 0, and we computed separate statistics for residues that were exposed versus buried in the protein interior (Fig. 3). The frequency of occurrence in the exposed outermost bins, between z = ± 34–40 Å was used as an aqueous standard state. The z-dependent profiles for the exposed positions provide an approximation of the free energy of transfer from water to various regions of the membrane (ΔG_PDB_) ^33,35^.

**Figure 3.**
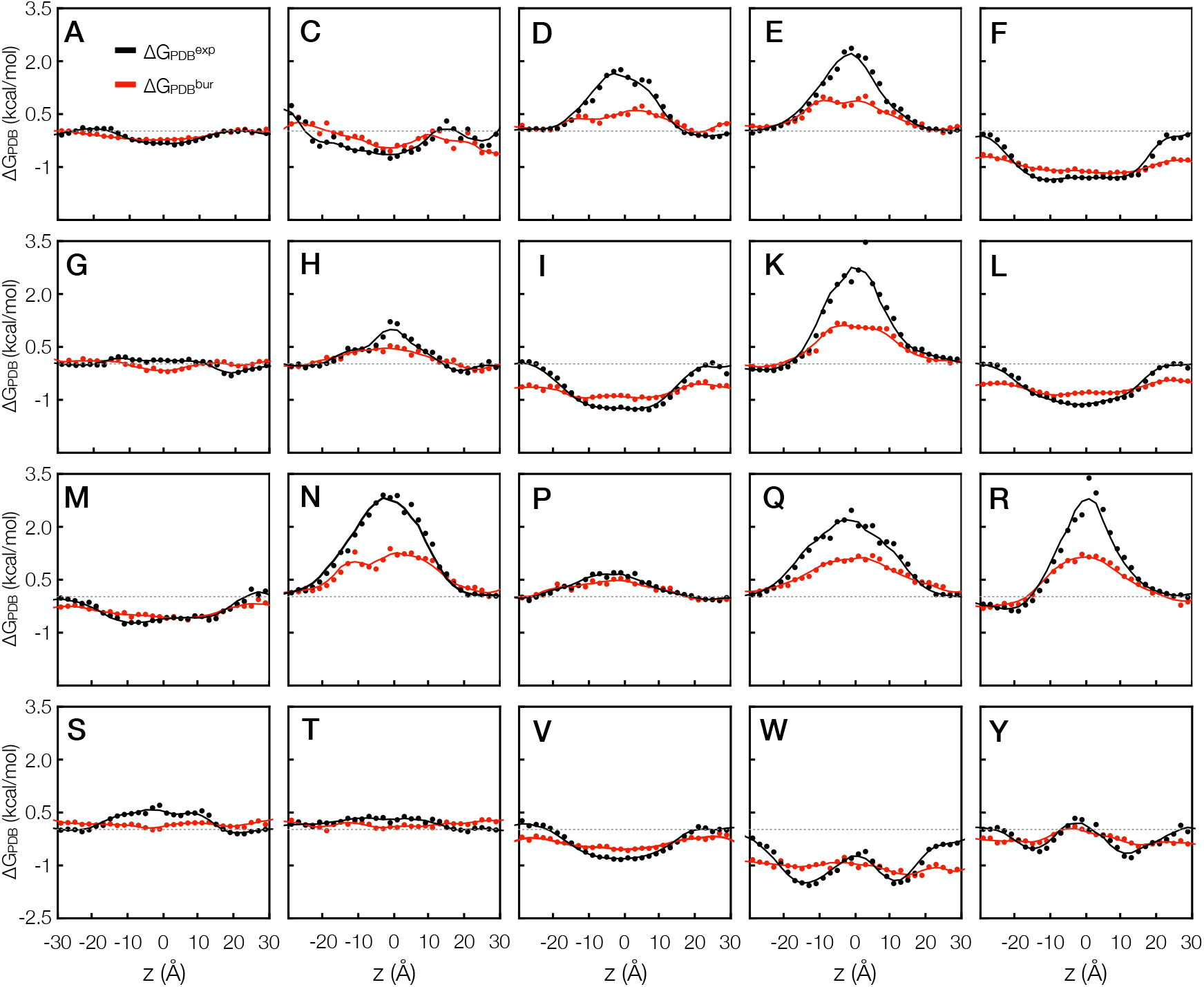
Free energy of transfer from water to various regions of the membrane (ΔG_PDB_) calculated from propensity differences. Each plot shows the free energy of transfer for moving an amino acid from an exposed position in water (averaged from propensities at z= ±34–40 Å) to a position at various z heights, either exposed at the protein exterior (ΔG_PDB_^exp^) or buried inside the protein (ΔG_PDB_^bur^). ΔG_PDB_ values are fitted with sigmoids, a Gaussian, or a combination of the two, as seen most appropriate (see Supplemental Table SX for parameters of fitted curves).

As expected, we find that the Z-dependent values of ΔG_PDB_ depend quite markedly on the burial of residues in a protein, with the exposed residues having a much greater tendency than buried residues to match their physical properties to that of the environment. Thus, the penalty for bringing a polar residue such as Asn, Asp, Gln, Arg, Lys to the center of the bilayer is 1 to 2 kcal/mol greater when their sidechains are lipid-exposed versus buried in the protein interior, where they can engage in favorable electrostatic and hydrogen-bonded interactions. Similarly, apolar residues have the strongest tendency to be exposed to the membrane lipids near the center of the bilayer. Finally, Trp and, to a lesser extent, Tyr have a pronounced tendency to accumulate at surface sites near the headgroup region, which display strong minima in ΔG_PDB_ at the headgroup region (approximately ± 12 to 15 Å relative to the bilayer center) for exposed but not buried positions. These findings help rationalize the stabilizing influence of Tyr and Trp on ΔG_app_^Wat→Int^ versus ΔG_app_^Wat→TM^ in our biological scales. Moreover, the aromatic residue Phe behaves more like simple apolar sidechains such as Ile and Leu than the interface-seeking Tyr and Trp residues in both the biological and PDB-derived scales.

To examine similarities among the amino acids, we clustered the 20 residue types based on their position-dependent values of ΔG_PDB_ (Fig. 4a). The ranking of the amino acids shows many expected features. For example, polar residues cluster together, with Lys + Arg and Glu + Asp in separate sub-clusters. Trp and Tyr cluster together, far from Phe, again showing that amphiphilicity plays an important role in defining locational preferences for these amino acids. Small polar (G, S, T) and hydrophobic residues (L, I, V, M, F, A) cluster together. The observed clustering is quite similar to the rankings seen in the values of ΔG_app_^Wat→Int^ and ΔG_app_^Wat→TM^.

**Figure 4.**
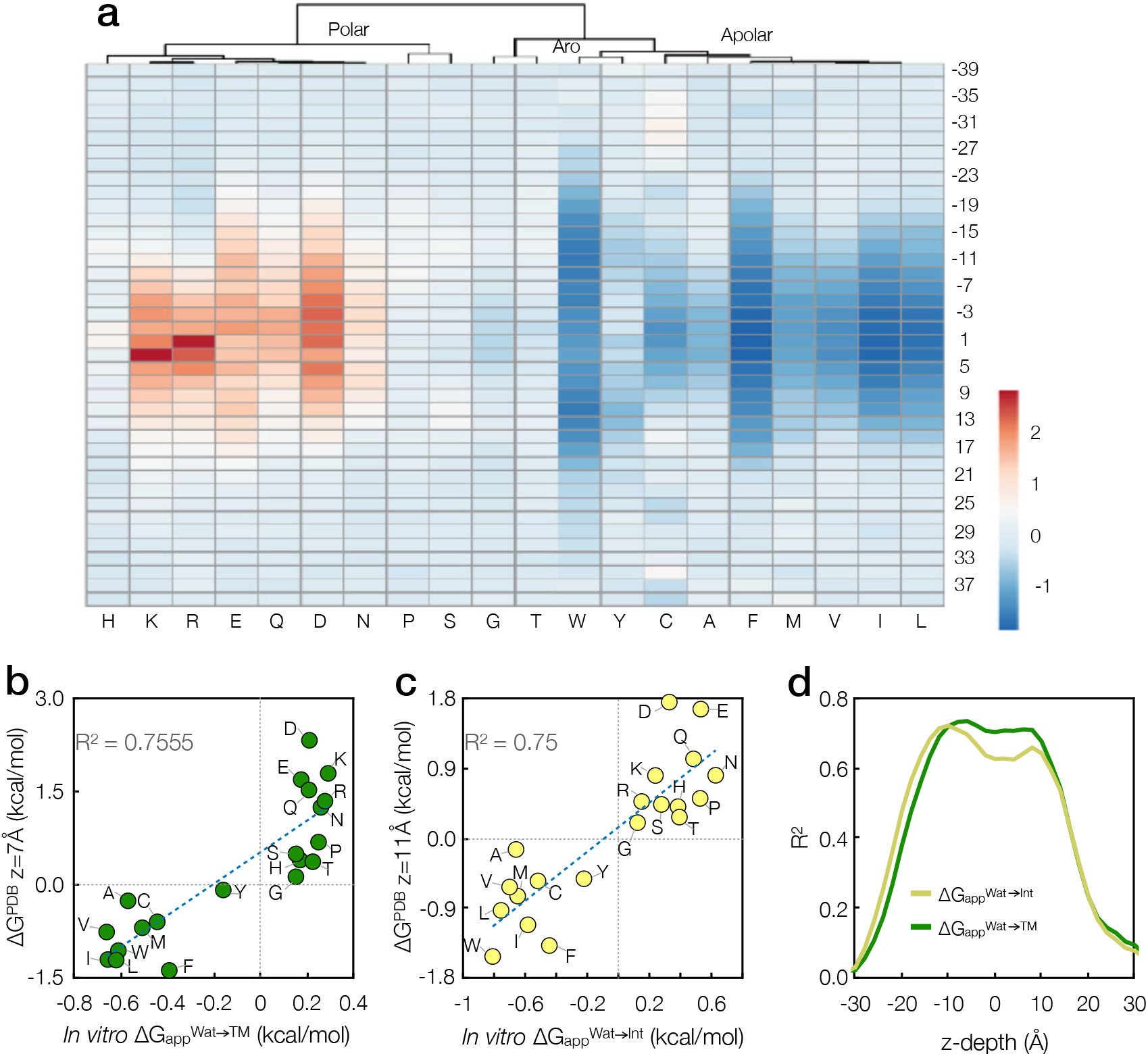
Comparison between experimentally derived scales and ΔG from propensity calculations. Amino acids are marked beside their corresponding data points by single-letter names. **a**. Heatmap-dendrogram of ΔG_PDB_ values at different z heights. Data are ΔG_PDB_ values of moving amino acids from a generic position in water to various positions in the membrane and exposed at the protein exterior. Data are clustered using the ClustVis web tool^47^. Columns are clustered using correlation distance and complete linkage. **b**. Experimentally derived water-to-TM scale (ΔG_app_^Wat→TM^) in the *in vitro* case compared to ΔG_PDB_ from an exposed position in water into a position exposed to the lipid bilayer at z = -7Å. **c**. Experimentally derived water-to-interface scale (ΔG_app_^Wat→Int^) in the *in vitro* case compared to ΔG_PDB_ from a generic position in water into a position exposed to the lipid bilayer at z = -11Å. **d**. Coefficient of determination (R^2^) values for linear least squares trendlines between all Z-dependent values of ΔG_PDB_ and ΔG_app_^Wat→TM^ (green line) and ΔG_app_^Wat→Int^ (yellow line) scale values.

We next compared our biological scales with the ΔG_PDB_ values computed at varying depths in the bilayer with the expectation that ΔG_app_^Wat→TM^ would correlate best with ΔG_PDB_ values computed near the bilayer center, while ΔG_app_^Wat→Int^ would correlate better with values computed near headgroup region. Indeed, ΔG_app_ ^Wat→TM^ correlates best with ΔG_PDB_ at z = -7 Å (Fig. 4b). This location is consistent with the position of the variable residues in the bRc peptide, which were slightly displaced from the TM region. By contrast, ΔG_app_^Wat→Int^ correlates best with ΔG_PDB_ at z = -11 Å (Fig. 4c). This value represents a minimum in ΔG_PDB_ for interfacially disposed residues such as Tyr and Trp. It also is a region where the value of ΔG_PDB_ is highly sensitive to z-position for both polar and apolar residues. Thus, even small adjustments in sidechain conformation or rigid-body shifts of an interfacial helix at this depth of insertion would cause a large change to the free energy of association. This heightened sensitivity manifests itself as a sharper peak in R^2^ for ΔG_app_^Wat→Int^ at z = -11 Å than the corresponding peak ΔG_app_^Wat→TM^ at z = -7 Å (Fig. 4d). Hence, these correlations are consistent with the behavior expected for the assumed equilibrium between TM and interfacial locations.

We also considered the possibility that differences in the experimental systems contribute to a lack of perfect correlation between the biologically and statistically derived scales. In particular, the interfacial helical state is not well represented in our PDB files, which are dominated by multi-span TM helices. We therefore sought reduced dimensionality representation of sequence space rather than structural space for water-soluble and membrane helices to allow comparison of our bRc peptide sequences to natural proteins.

### UMAP analysis of helical peptide segments

Uniform manifold approximation and projection (UMAP) has recently emerged as a useful means of visualizing high-dimensional data in a low-dimensional representation ^36^. To better understand how our experimentally characterized sequences relate to the broader natural distribution of helical protein segments, we generated a UMAP from a dataset of 3,672 sequences of helices from soluble proteins and 2,941 sequences of TM helices from the Uniprot database^37^. Each sequence was converted into a 24-feature vector consisting of the fractional amino acid composition, the average of the z-dependent ΔG_PDB_ values computed for each amino acid by assuming the helix spans the membrane (Materials and Methods), the hydrophobic moment computed using the same ΔG_PDB_ values, the net charge of the peptide segment at pH 7, and the length of the sequence.

The resulting UMAP shows a two-lobed structure (Fig. 5a) with sequences from soluble helices in one large cluster (left lobe) and sequences from TM helices in the other (right lobe), with minimal overlap. When the bRc derived sequences studied here are embedded into the same UMAP their locations within the two lobes were reflective of the propensities of the helices for the aqueous, interfacial, and TM states. The individual points are aggregated into eight clusters (Supplementary Table 2), to allow us to examine how a position in the UMAP relates to the f_Wat_, f_Int_ and f_TM_. Satisfyingly, there is qualitative agreement between the positions in the UMAP versus the fractions of these states, with the TM and Interfacial states being increasingly populated as one moves from left to right. To place this qualitative observation on more quantitative footing, we derived moments describing the predilection of the helices to form the three species based on their position in the UMAP (Fig. 5a). The three moments represent a direction along which the embeddings are most highly correlated with the respective probabilities of the three states. The results show good agreement with the qualitative observation that the horizontal roughly determines partitioning between water and TM states. The vertical axis reflects additional information relating to increased interfacial propensity.

**Figure 5.**
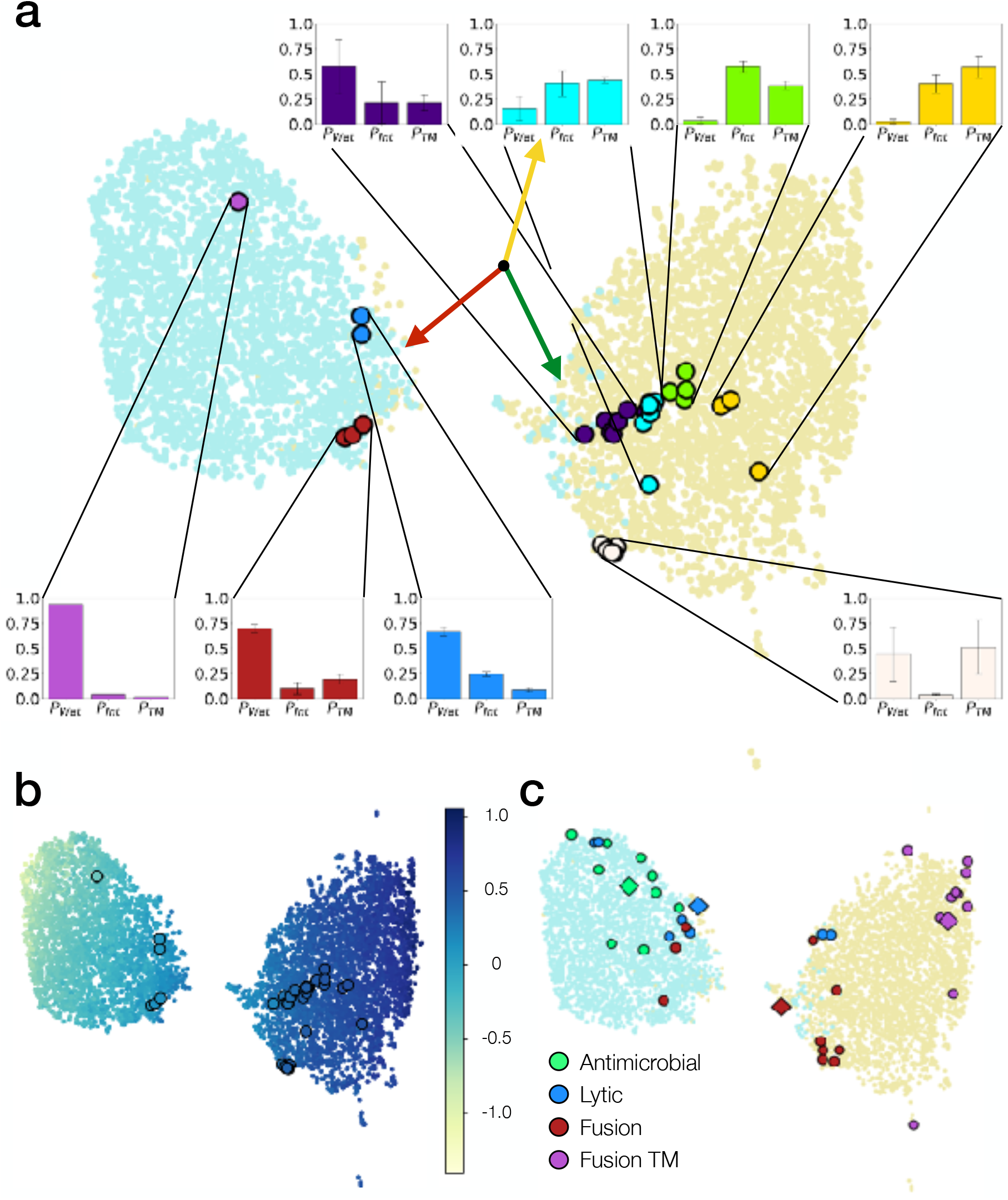
UMAP embeddings of peptide sequences from Uniprot exhibiting known helical disposition within soluble or transmembrane proteins. The experimentally tested sequences are embedded in the same space as large points with black outlines. **a**. UMAP embeddings with Uniprot-derived sequences colored in light blue if found in soluble domains and beige if found in transmembrane domains. The UMAP embeddings cluster into two large lobes, largely delineating helices within soluble proteins from transmembrane helices. All sequences tested with the LepB glycosylation assay (Supplementary Table 2) were visually clustered based upon their embeddings and the predicted statistics over the three dispositions (aqueous (*P*_*Wat*_), interfacial (*P*_*Int*_), and transmembrane (*P*_*TM*_)) are shown as histograms. Each of the three helical dispositions rapidly becomes more dominant along a particular direction in the UMAP space, shown by the three vectors utilizing the same color scheme for the three dispositions as in Fig. 1d (green, yellow, and red for inserted, interfacial and translocated location, respectively). **b**. The same UMAP embeddings, colored according to average hydrophobicity computed using the ΔG_PDB_ scale, show a clear trend toward higher hydrophobicity (i.e. lower average ΔG_PDB_) for helices from soluble proteins from left to right. **c**. UMAP embeddings of helical peptides with known function ^20^ appear in distinct regions of the UMAP space. Antimicrobial peptides, which tend to be soluble and negatively-charged, appropriately cluster in the lower upper half of the left lobe. Lytic peptides and the C-terminal segments of viral fusion proteins occupy the boundary region within which both helices from soluble proteins and transmembrane proteins can be found. Transmembrane viral fusion domains, which must insert into a transmembrane state to function, can be found at the rightmost edge of the right lobe, implying a strong preference for the transmembrane disposition. Points labeled with circular markers represent individual sequences, while diamonds represent means in the UMAP space over a functional family.

Lastly, we analyzed the capacity of the UMAP projection to meaningfully cluster the sequences based upon physical properties and functional families. Coloring the embeddings by physical properties, we found the horizontal axis of the UMAP reflects hydrophobicity as assessed by our lipid-exposed ΔG_PDB_ scale (Fig. 5b), while the vertical axis reflects net charge at pH 7 within the left lobe (Supplementary Fig. 10a). Sequences with high helical hydrophobic moments, also calculated using our lipid-exposed ΔG_PDB_ scale, congregated in the upper left section of the left lobe (Supplementary Fig. 10b). To examine whether the UMAP could meaningfully cluster peptide sequences with different functions, we embedded the membrane-associating sequences from four distinct functional classes of previously characterized peptides and proteins whose functions are dictated by their ability to bind to membrane surfaces into the UMAP (Fig. 5c).

We first examined helical antimicrobial peptides (AMPs) and cytotoxic peptide sequences. Both bind to membrane surfaces, attracted to microbial bilayers that are richer in acidic phospholipids than mammalian membranes, while cytotoxic peptides bind more indiscriminately to eukaryotic and bacterial membranes. Both disrupt cell membranes by mechanisms that involve surface-absorbed as well as membrane-spanning states. Our analysis shows that AMPs embed into the basic and water-soluble helical regions of the UMAP, while cytotoxic peptides tend to have embeddings directly between the TM and water-soluble states. Thus, on average cytotoxic peptides would be predicted to have higher affinities for membranes and be more stable in TM states. This finding is in keeping with their more aggressive and non-selective behavior towards bilayers relative to AMPs. (For references on this subject see the review ^31^ and other reviews referenced within).

Helical fusogenic sequences present another interesting class of proteins, which are released from membrane-embedded fusogenic proteins to bind to the surface of target membranes and ultimately help stabilize a dynamic membrane-spanning fusion pore. The dichotomy between the embeddings for the TM regions of fusogenic peptides and their fusion pores is functionally interesting in this regard. The TM regions lie in the far right of the plot, consistent with their function as membrane anchors, while the fusogenic peptides lie close to the interface between water-soluble and membrane-spanning helices in keeping with their dynamic requirements for membrane binding. These considerations extend our analysis of model peptides to sequences that, like the bRc peptides, are poised for membrane insertion and surface association.

## Discussion

This work provides the first large-scale, systematic examination of the role of sidechains in mediating the propensity of peptides in a helical conformation to associate with the membrane interface during translocon-mediated protein insertion into the endoplasmic reticulum. In pioneering biophysical studies ^18,38^, Wimley, White and coworkers examined short (non-helical) host-guest pentapeptides to see how variations in sequence led to changes in the energetics of binding to the surface of artificial liposomes to provide the first experimental scale for surface association. Their interfacial scale showed a significant correlation with hydrophobic scales, such as one derived from octanol-water partition coefficients (R^2^ = 0.86) (Supplementary Fig. S11). Amphiphilic aromatic residues, Trp and Tyr, showed enhanced affinity for the membrane interface, although charged residues such as Arg and Lys, which clearly contribute to interfacial recognition in our own and previous studies ^39^, did not appear to enhance affinity for the neutral lipid bilayers used in these studies. We also observe good correlation between ΔG_app_^Wat→Int^ and other hydrophobicity scales (Supplementary Fig. S12), most notably the z-dependent scale for inserted helices elucidated by Hessa et al. (Fig. 2b, 2c).

Sequence-based bioinformatic analysis via UMAP also provided complementary insights to the experimental assay. The clustering of sequences in the UMAP space based upon aqueous or transmembrane disposition, as well as net charge, hydrophobicity, and known function, showed that UMAP could embed short peptide sequences in semantically meaningful ways. Each experimentally-tested sequence had numerous close neighbors from endogenous proteins, representing a broad sample of the known sequence space. Moreover, they clustered near other experimentally tested sequences with similar propensities for the three states, with the horizontal coordinate in the UMAP space roughly determining the partitioning between the aqueous and TM states and, for more hydrophobic sequences, the vertical coordinate roughly determining the partitioning between the aqueous and TM states. We hypothesize that endogenous sequences near to the experimentally tested sequences in the UMAP space will have comparable preferences for the three states, presenting a potential direction for future study.

Finally, both the experimental ΔG_app_^Wat→Int^ and the informatic scale, ΔG_PDB_, provide important information for the *de novo* design of membrane proteins. The large difference between ΔG_PDB_ for buried versus exposed sites has not previously been appreciated. The scale was derived in a manner that allows easy incorporation into algorithms for sequence design given a backbone structure. Thus, our studies should enable understanding of natural proteins as well as *de novo* design of novel peptides and proteins in the highly heterogenic milieu of the membrane, spanning a wide range of topologies and functions. We posit, on the basis of our findings, that the vital process by which cells determine the relation of their myriad proteins to the plasma membrane can be reduced to a collection of straightforward sequence-based rules.

## Online Methods

### Enzymes and chemicals

All enzymes as well as plasmid pGEM1 and the TnT coupled transcription/translation system were from Promega (Madison, WI). SP6 RNA polymerase and ER rough microsomes from dog pancreas were from tRNA Probes (College Station, TX). EasyTagTM EXPRESS35S Protein Labeling Mix, [35S]-L-methionine and [35S]-L-cysteine, for *in vitro* labeling was purchased from Perkin Elmer (Waltham, MA, USA). Restriction enzymes and Endoglycosidase H were from Roche Molecular Biochemicals (Basel, Switzerland). The DNA plasmid, RNA cleanup, and PCR purification kits were from Qiagen (Hilden, Germany). The PCR mutagenesis kit QuikChange was from Stratagene (La Jolla, CA). All the oligonucleotides were from Macrogen Inc. (South Korea).

### DNA Manipulation

Oligonucleotides encoding the different bRc variants and melittin flanked by GGPG…GPGG tetrapeptides intended to “insulate” the central peptide from the surrounding LepB sequence were introduced into the pGEM1Lep plasmid^15,40^ between the *Spe*I and *Kpn*I sites by using four double-stranded oligonucleotides (38-58 nucleotides long) with overlapping overhangs at the ends and phosphorylated at 5 ’ ends. Pairs of complementary oligonucleotides were first annealed at 85ºC for 10 min followed by slow cooling to 30 ºC. After that, the pair-annealed double-stranded oligos were mixed, incubated at 65 ºC for 5 min, cooled slowly to room temperature, and ligated into the vector. All bRc inserts were confirmed by sequencing of plasmid DNA (Macrogen).

The bRc site-directed mutagenesis was performed using the QuikChange mutagenesis kit (Stratagene) following the manufacturer’s protocol. The DNA encoding LepG3 proteins was PCR amplified to incorporate a c-Myc tag at the N-terminus and subcloned into the mammalian pCAGGS vector for *in vivo* assays using the In-Fusion HD technology (Takara), according to the manufacturer’s instructions. All DNA manipulations were confirmed by sequencing of plasmid DNAs. Site-directed mutagenesis was also used to introduce acceptor sites for N-linked glycosylation at appropriate positions.

### *In vitro* transcription and translation

LepB-bRc constructs were transcribed and translated in the TnT Quick system (Promega). 1 μg DNA template, 1 μL (5 μCi) ^35^S-Met/Cys (PerkinElmer) and 1 μL microsomes (tRNA Probes) were added at the start of the reaction, and samples were incubated for 90 min at 30º C. After polypeptide synthesis membranes were collected by ultracentrifugation and analyzed by SDS-PAGE, gels were finally visualized on a Fuji FLA3000 phosphorimager using the Image Gauge software.

For the proteinase K protection assay, the translation mixture was supplemented with 1 μL of 50 mM CaCl2 and 1 μL of proteinase K (4 mg/mL) and then digested for 20 min on ice. Adding 1 mM PMSF before SDS-PAGE analysis stopped the reaction.

After polypeptide synthesis membranes were collected by ultracentrifugation and analyzed by SDS-PAGE, gels were finally visualized on a Fuji FLA3000 phosphorimager using the Image Gauge software.

### Mammalian cells assay

Hek293T cells were grown in Dulbecco’s modified Eagle’s medium (DMEM) (Gibco) supplemented with 10% fetal bovine serum (FBS), and penicillin-streptomycin (P/S) (100 U/mL) at 37 ºC, 5% CO_2_. Cells were plated in 24-well plates (2×10^6^ cells/plate) and transfected after 24h. For transfection procedure, 500 ng/well of plasmids encoding c-Myc tagged LepG3 were added to a mixture of 2 µL PEI MW 25,000 (1 mg/mL) (Alfa Aesar) diluted in 100 µL Opti-MEM reduced serum medium (Gibco). Transfection mixture was incubated for 20 min at RT and then added dropwise to 24 h cultured cells in 500 µL of DMEM containing FBS and P/S.

At 48 h post-transfection cells were collected in lysis buffer (TBS (Tris-HCl 20 mM pH7.5, NaCl 150 mM), 1% SDS) and put through three freeze-thaw cycles. The suspensions were clarified by centrifugation (13,000 x g). Supernatants were mixed with SDS-PAGE sample buffer, heated 5 min at 95 ºC, and loaded on 12% SDS-PAGE. Next, proteins were transferred onto PVDF membranes. Immuno-identification of the LepG3 system proteins was done using α-c-Myc antibody (Merck) followed by a secondary HRP-conjugated α-rabbit antibody (Merck). Chemiluminescence was visualized by an ImageQuant LAS 4000 (GE Healthcare). Bands were quantified using ImageJ (NIH).

### Peptide synthesis

The bRc peptides were synthesized using standard Fmoc chemistry by automated microwave-assisted solid phase peptide synthesizer (Biotage Initiator+Alstra) on 0.1 mmol scale. Each cycle included (1) Fmoc deprotection (20% 4-methyl piperidine with HOBt (0.1 M) in DMF, 4.5 mL, 5 min, 75 °C); (2) Coupling with N-α-Fmoc-amino acid (5 eq, 0.5 M in DMF), HCTU (4.95 eq, 0.5 M in DMF), DIEA (10 eq, 0.5 M in DMF) (5 min, 75 °C). N-terminal acetylation was done by treatment or resin with Ac_2_O (10 eq) and DIPEA (20 eq) in DMF and the final cleavage was performed using TFA:TIPS:H_2_O (95:5:5). The crude peptide was obtained by cold ether precipitation and purified by RP-HPLC. Their chemical entity and purity were confirmed by MALDI and analytical HPLC.

### CD spectrometry

Small unilamellar vesicles (SUVs) of POPC were prepared by combining ethanolic solution of POPC and the peptide ([peptide]/[lipid] = 1/100), drying to a film and lyophilized for overnight. The thin film was resuspended in phosphate buffer (20 mM, pH = 8.0) to a phosholipid concentration of 25 mM and tip-sonicated (Fisher Sonic Dismembrator Model 500, 20% power, 5 min, 2 s on, 2 s off). The resulting mixture was further diluted to 5 μM (peptide concentration) in 3 mL of phosphate buffer (20 mM, pH = 8.0) in a 1-cm cuvette. Circular dichroism data were collected on a JASCo J-810 (8 s average, 2nm bandwidth, triplicate measurement).

### Analysis of the observed free energy of transfer from the experimental data

The fraction of the peptide sequences in the water-exposed, interfacial, or TM locational states were approximated from their relative mole fractions. To obtain these mole fractions from the experimentally observed radiometric counts, a statistical model was developed to account for the effects of incomplete glycosylation of the nascent chain. Beginning with the counts *C*_*i*_, with *i* ∈ {1,2,3} denoting the number of glycosylations, the probabilities *P*(*i*) of each observed number of glycosylations were calculated as:

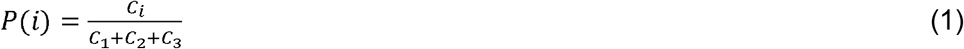

The glycosylation state probabilities of Eq. 1 were then expressed in terms of the conditional probabilities of observing *i* glycosylations given that the peptide sequence is in a known locational state, as follows:

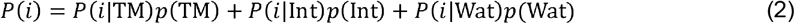

The three instances of Eq. 2, one for each, may be collected into a matrix-vector product:

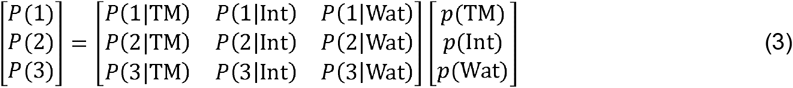

The columns of the matrix may be interpreted as the probabilities of observing each of the three possible glycosylation states given that the protein is in a known locational state. While the TM state is assumed to never be doubly or triply glycosylated and the interfacial state is assumed to never be triply glycosylated, incomplete glycosylation by the OST is expected to lead to a mixture of single and double glycosylation for the interfacial state and single, double, and triple glycosylation for the aqueous state. For *P*(2|Int), we retained to three significant digits the baseline double glycosylation probability of 86.5% from the original study by Hessa et al. ^15^, while the values of *P*(*i*|Wat) were determined from our experimental glycosylation data for the soluble sequence GDKQEGEWPTGLRLSIGGI (corresponding to residues 304-322 in the translocated P2 domain of LepB). We computed *P*(*i*|Wat) for *i* = 1,2,3 by averaging the glycosylation state probability *P*(*i*), determined via Eq. 1, across the four experimental replicates for the P2 domain. This analysis generated the following values:

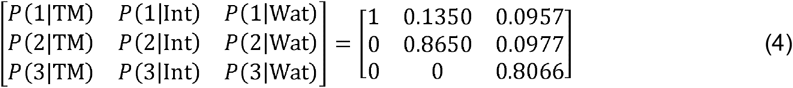

As the matrix given in Eq. 4 is invertible, the locational state probabilities were determined from multiplying both sides of Eq. 3 by the inverse matrix. As a result:

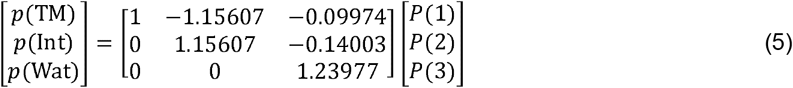

Locational state probabilities were estimated from the glycosylation state probabilities via Eq. 5 for all peptide sequences that were experimentally tested. To amplify the apparent differences in mole fraction between the singly- and doubly-glycosylated states, three substitutions were simultaneously made, replacing a DLP sequence (residues 11, 12 and 13 in bRc-derived sequence) with a new sequence X_1_X_2_X_3_. Free energy differences between the water-exposed state and the interfacial and TM states, respectively, were calculated from probability ratios as follows:

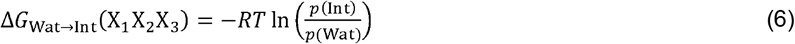

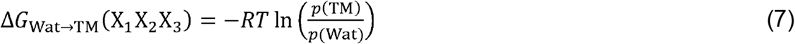

In Eqs. 6 and 7, the value of 0.602 or 0.612 kcal mol^-1^ was used for the ideal gas constant and temperature, *RT*, for *in vitro* or *in vivo* data respectively. The dependence of these free energy differences upon the sequence X_1_X_2_X_3_ is modeled as a sum over ΔΔ*G* values associated with the substitution Z_i_ → X_i_, wher Z_1_Z_2_Z_3_ is a hypothetical sequence for which Δ*G*_Wat→Int_ (Z_1_Z_2_Z_3_) = 0 and Δ*G*_Wat→TM_ (Z_1_Z_2_Z_3_) = 0. Formally:

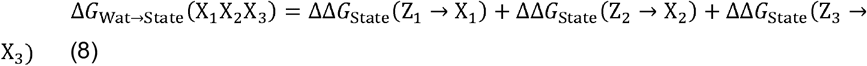

While the form of Eq. 8 implies a positional dependence of Δ*G* upon the three substitutions, we saw that permutations of the sequences AAG and AGG showed much smaller differences in Δ*G*_Wat→TH_ or Δ*G*_Wat→Int_ between sequences of the same composition versus between sequences of different composition (Supplementary Table S3). Thus, we assume Δ*G* is independent of order and thus 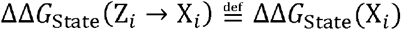. Therefore, Δ*G* can be expressed as a sum over all 20 amino acids X of the corresponding ΔΔ*G*_state_ (X_*i*_) values, weighted by the number of occurrences *n*_*X*_ of the amino acid in the sequence:

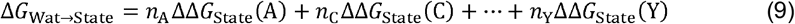

A total of 113 replicates were tested experimentally across 27 sequences X_1_X_2_X_3_, including DLP and the 20 possible sequences XXX for each of the amino acids. The replicates for each sequence were analyzed for outliers that deviated from the sample mean by more than twice the sample standard deviation, resulting in the identification of one outlier for the sequence AAA, which was discarded. The 112 remaining instances of Eq. 9 for each of the experiments were concatenated into matrix form:

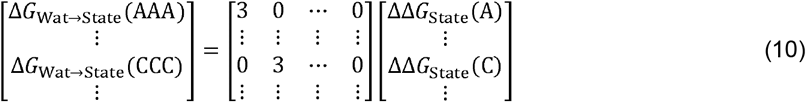

The two vectors of 20 values for ΔΔ*G*_Int_(X) and ΔΔ*G*_TM_(X) were estimated by weighted linear least-squares regression using the NumPy Python package^41^. The weights were set to be the sample standard deviations of the Δ*G*_Wat→State_ values across the experimental replicates of each three-residue sequence. The errors are given by the standard errors of each regression variable.

### Database for Propensity Calculation

Distinct structures of bitopic and alpha-helical polytopic proteins from the OPM database^34^ as of Feb. 2022 were used for propensity calculation. Proteins in this database are in their native biological assemblies, have an average TM helix tilt of 0°, and the average centroid of the TM region is at the bilayer center. Therefore, their structures were used without further alignment. Only structures, or chains in multimeric structures, with resolution better than 3.5 Å and a maximum sequence identity of 70% were selected for further analysis. The final list of PDB accession codes and chain IDs for propensity calculation can be found in **Data S1**. The list contains a total of 2,193 structures with 7,057 unique chains. Note that biological assemblies were used for the correct identification of interior(buried)/exterior(exposed) residue positions, while the asymmetric units of the structures from this nonredundant list were used for all other statistics and calculations.

### Propensity Calculation

Propensities were calculated in a similar fashion to previous results^20,34,42^. Structures from the OPM database were aligned so that the membrane normal was parallel to the z-axis, with z = 0 at the bilayer center and negative z toward the cytoplasm. Residue positions were defined by the coordinates of C_β_ (C_α_ for Gly). The structural data was divided into bins of 2 Å along the z-axis and the occurrence of each residue at each binned z-value was counted. Propensities were calculated using the equation:

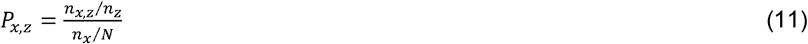

where *P*_*x, z*_ is the propensity of a given residue *x* in a given z-value bin *z, n*_*x, z*_ count of residue *x* in bin *z, n*_*z*_ is the total count of all residue types in this bin, *n*_*x*_ is the total count of residue *x* in all bins, and *N* is the total count of all residue types in all bins.

Residues were identified as “exterior” (exposed) or “interior” (buried) using a convex hull algorithm^43,44^. The coordinates for C_α_ and C_β_ atoms were used to define two surfaces. If a C_β_ atom fell outside the surface of the C_β_ hull, that residue was identified as “exterior”; conversely, if a C_β_ atom was found within the surface of the C_α_ hull, that residue was identified as “interior” (C_β_ atoms were appended to the C_α_ atoms of Gly for this calculation by converting the residue to Ala). The advantage of using C_α_ and C_β_ atoms is that this treatment can be used in future studies for protein design, in which the backbone but sequence has been computed (in which case the values of ΔΔG_PDB_ can be used directly in a design energy function or to bias sampling). Moreover, this approach eliminates problems with missing sidechains or limited resolution for surface residues in experimental structures. The radius of the alpha-sphere used to define the surfaces was 8 Å. Propensities for “exterior” (exposed) and “interior” (buried) residues were calculated the same way as described above, but for “exterior” and “interior” residues, respectively.

### ΔG_PDB_ calculation

For the “reaction” of moving a residue from one position (e.g., in the aqueous environment) to another position (e.g., in the interior of a protein at z = 0 Å), a pseudo free energy ΔG_PDB_ was defined by Eq. 12:

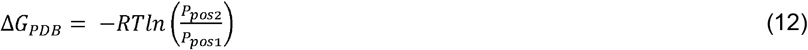

where *P*_*pos*1_ and *P*_*pos*2_ are the propensities of this residue at positions 1 and 2, respectively, is the gas constant, and is temperature in Kelvin. As in Equations (6) and (7), the value of 0.593 kcal mol^-1^ was used for the ideal gas constant and temperature, *RT*, in Equation (12). Considering that a residue is sufficiently distant from the lipid membrane when z < -35 Å or z > +35 Å, the propensity for a residue to occupy the aqueous environment was defined by the average of its propensities at the two z-value ranges of -39 to -35 Å and +35 to +39 Å.

### UMAP Analysis

Protein sequences of known α-helices were retrieved from the Uniprot database^37^. The sequences corresponding to TM helices were derived from integral membrane proteins with no more than four membrane-spanning segments. This was done to ensure that all sampled helices are at least partially exposed to the lipid environment and not buried within the cores of large transmembrane protein domains. Additionally, given that TM helices must span the bilayer and thus are seldom shorter than 20 amino acids in length, a minimum length of 15 amino acids was imposed upon the helices from soluble proteins. This resulted in a dataset consisting of 3,672 transmembrane helices and 2,941 helices from soluble proteins.

Each α-helix was featurized as a 24-dimensional vector, with entries given by the fractional composition of the sequence by each of the 20 amino acids, the average z-dependent ΔG_PDB_ across the sequence, the hydrophobic moment computed using the calculated ΔG_PDB_ values for each residue, the net charge *Q*_net_ at pH 7, and the length *L* of the peptide segment. The z-dependent ΔG_PDB_ values were calculated according to Equation (12) under the assumption that the TM reference state consists of the apolar portion of a membrane helix inserted at an angle across the membrane with its centroid at z = 0 and the angle determined such that the N- and C-termini are at z = 15 and z = -15, respectively. Helices with *L* ≤ 21 are assumed to be oriented perpendicular to the bilayer normal. Formally, for residue indices *i* ∈ ⟦1, *L*⟧:

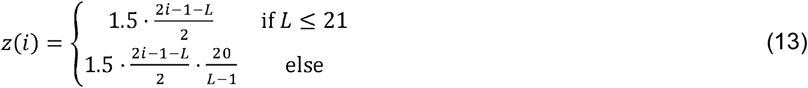

For *L* > 21, Equation (13) linearly interpolates the interval [−15, 15] to determine the z-coordinates of each residue, which is equivalent to averaging the z-coordinate of each residue over all possible rotations of the helix about its axis (Supplementary Fig. 12). Hydrophobic moments were computed using the method of Eisenberg et al.^45^ Moreover, because the net charge at pH 7 and helix length were initially much larger in magnitude than the other 22 features, they were normalized to lie within the interval [0, 1] as follows;

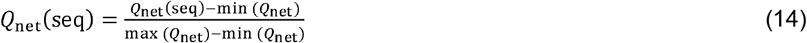

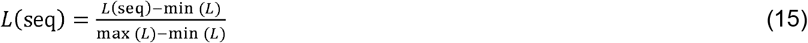

The 24-dimensional feature vectors from the Uniprot-derived database were then passed as input to the UMAP fitter in the UMAP Python package, using default parameters. Feature vectors for the experimentally-tested sequences and the sequences from the four functional peptide classes were preprocessed using the minimum and maximum values for *Q*_net_ and from the Uniprot-derived database and embedded in the previously-fit UMAP space for comparative analysis.

## Abbreviations

ER: endoplasmic reticulum
LepB: leader peptidase
SDS-PAGE: sodium dodecylsulfate polyacrylamide gel electrophoresis
TM: transmembrane
UMAP: uniform manifold approximation and projection.

## Acknowledgments

This work was supported by grants PID2020-119111GB-I00, PID2023-152568NB-I00 and PRX21/00348 from the Spanish Ministry of Science, Innovation and Universities (co-financed by European Regional Development Fund of the European Union) and CIPROM/2022/062 from the Generalitat Valenciana (to I.M.). R.C.K. was supported by the U.S. Department of Defense (DoD) through the National Defense Science & Engineering Graduate (NDSEG) Fellowship Program.

