## Supplemental figures for "Sequence-dependent scale for translocon-mediated insertion of interfacial helices in membranes"

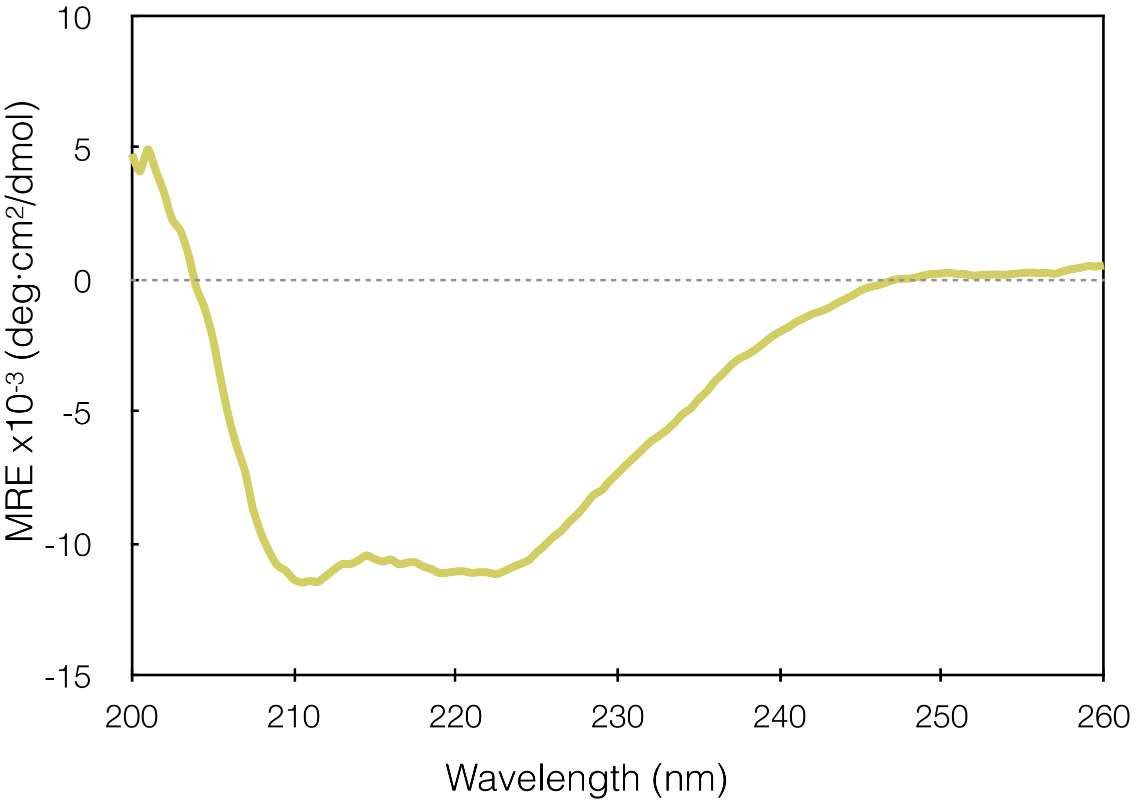


**Figure S1.** CD spectra of the bRc-derived peptide in the presence of POPC liposomes.

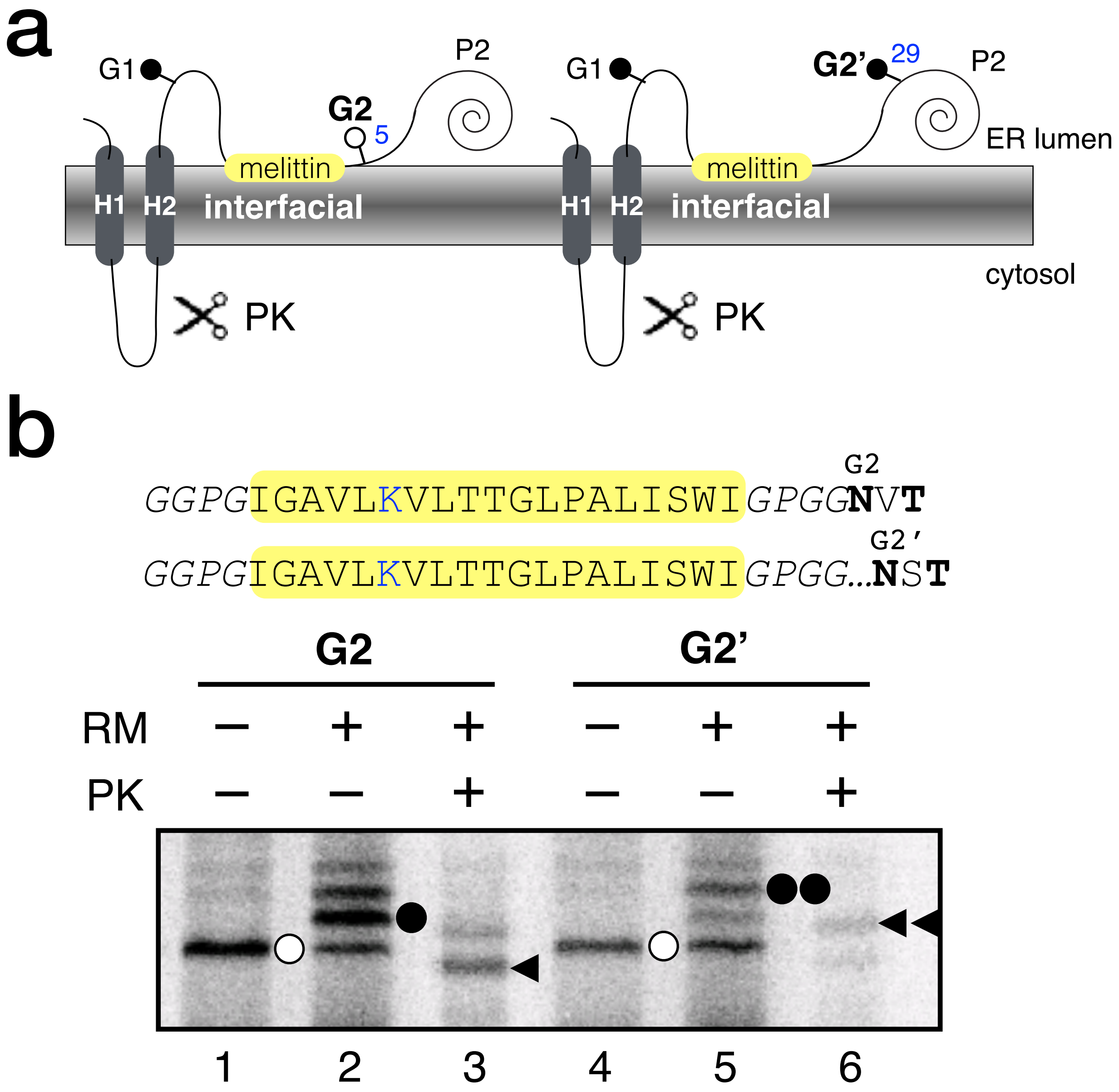


**Figure S2.** Interfacial disposition of melittin in the microsomal membrane. **a** wild type Lep has two N-terminal TM segments (H1 and H2) and a large luminal domain (P2) in which interfacial segment was inserted. An N-terminal glycosylation acceptor site (G1) was placed in positions 96-98. C-terminal glycosylation acceptor sites were placed next to the insulating GPGG sequence (G2) or 29 residues C-terminal from the insulating sequence (G2'). **b** Plasmids encoding the Lep/mellittin constructs were transcribed and translated in vitro in the presence (+) and absence (-) of dog pancreas rough microsomes (RM) and proteinase K (PK). Tested sequences are highlighted in a yellow box. Bands of non-glycosylated protein are indicated by a white dot; singly and doubly glycosylated proteins are indicated by one or two black dots, respectively. The arrowhead identifies undigested protein after PK treatment. One and two arrowheads indicate singly and doubly glycosylated protected fragments, respectively.

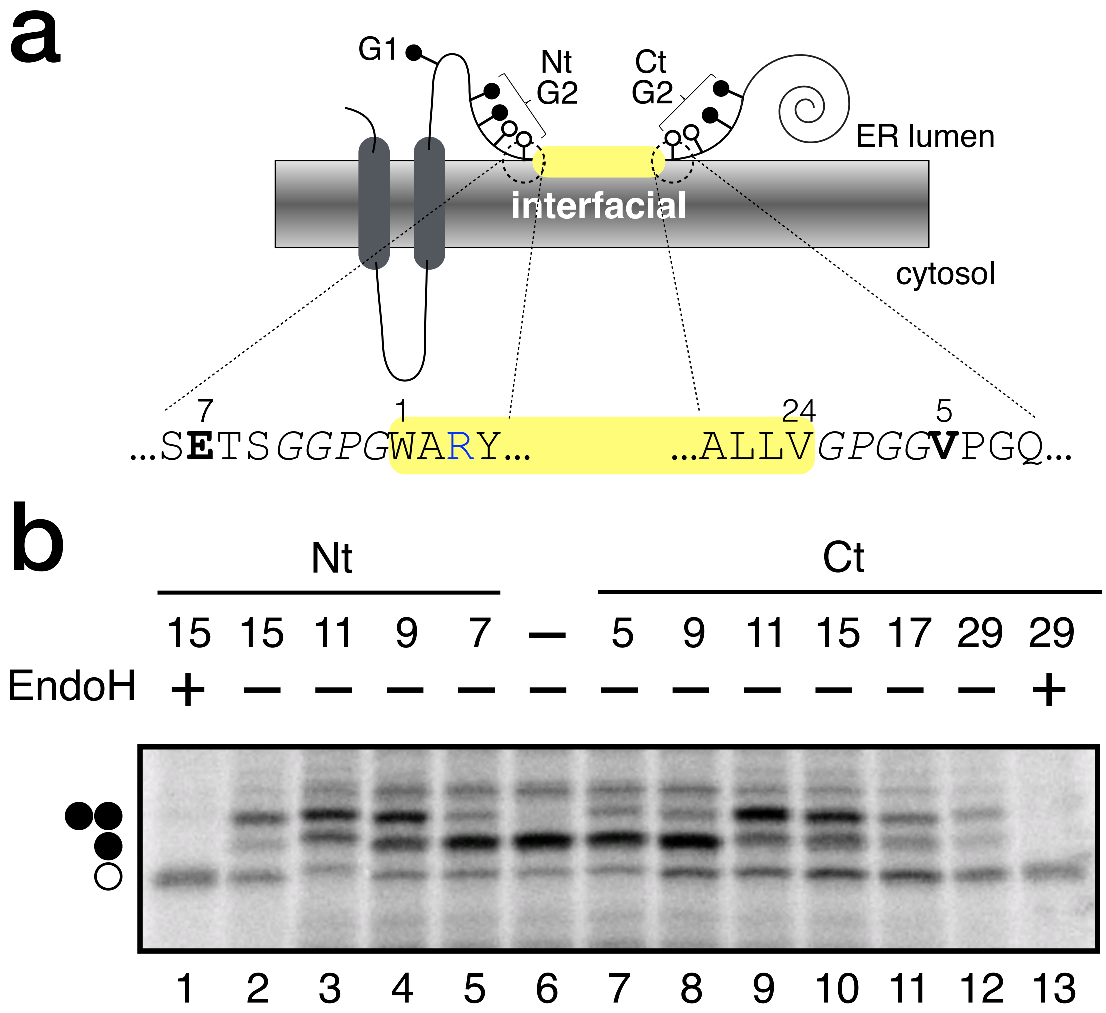


**Figure S3.** N- and C-termini minimal glycosylation distance for interfacial helices. **a,** Schematic representation of the LepG2 vehicle used. bRc derived sequence was flanked by appropriate glycosylation acceptor sites (see Table S1 for description of the mutants). **b**, In vitro translation of constructs with the G2 site at different amino acid distances from both ends of the interfacial sequence. Singly and doubly glycosylated forms of the protein are indicated with one or two black dots, respectively. The construct containing only G1 site (–, lane 6) was used as a control. Endoglycosidase H (EndoH) treatments were used to determine the non-glycosylated form of the proteins (white dot). **c**, Glycosylation profile quantification for constructs with the indicated distances between the interfacial bRc sequence and the G2 glycosylation acceptor site. Error bars represent the mean ± SD; obtained from at least 3 independent replicates.

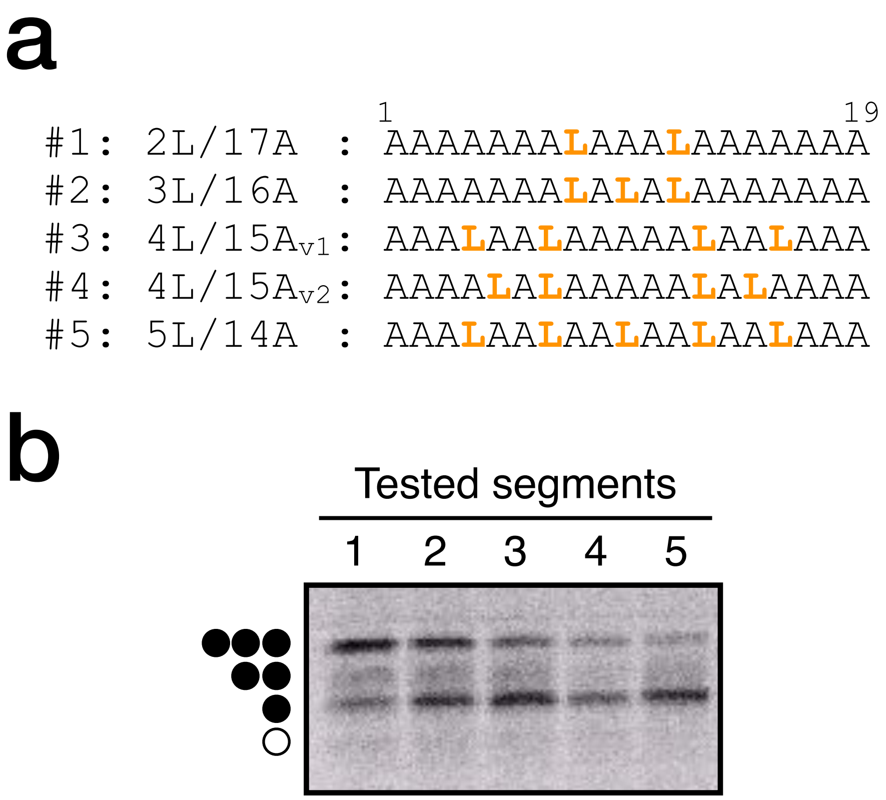


**Figure S4.** Membrane integration of tested segments with Leu/Ala composition 2L/17A, 3L/16A, 4L/15A, 5L/14A. **a**, Sequences of increasing number of Leu assayed in an Ala backbone. Incorporated Leu residues are shown in orange. **b**, Representative gel of the in vitro translation of the different constructs in the presence of microsomal membranes. Bands of non-glycosylated protein are indicated by a white dot; single, double, and triple glycosylated proteins are indicated by one, two, or three black dots, respectively.

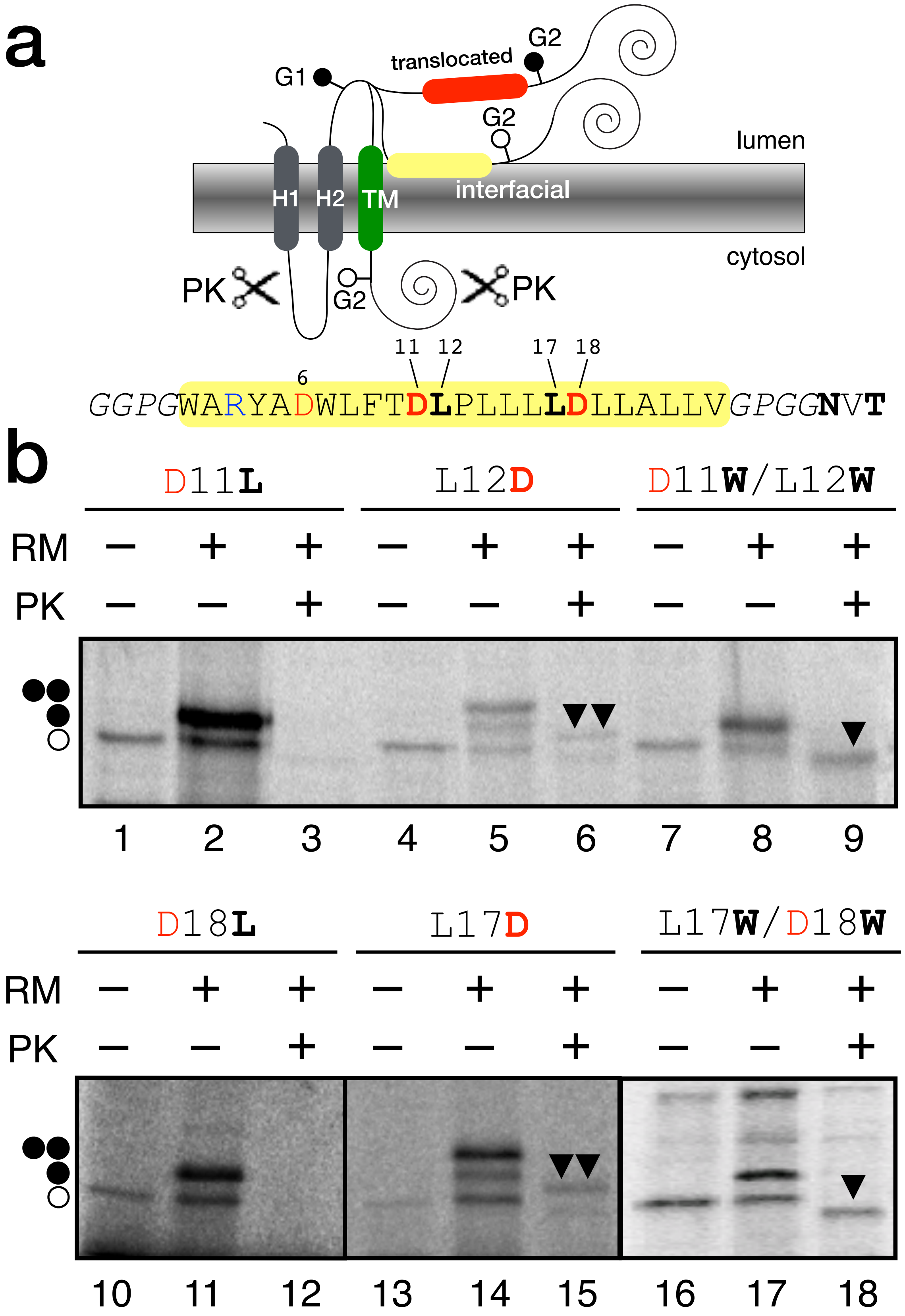


**Figure S5. bRc derived sequence screening for systematic substitutions. a,** Schematic representation of the LepG2 vehicle. bRc derived sequence is shown at the bottom and yellow boxed (pair residues tested highlighted in bold). **b**, bRc derived constructs translated in the absence (—) and in the presence (+) of rough microsomes (RM) and/or proteinase K (PK), containing single-point mutants D11L, L12D, D18L and L17D (lanes 1-3, 4-6, 10-12 and 13-15, respectively) and double mutants D11W/L12W and L17W/D18W (lanes 7-9 and 16-18, respectively). Bands of non-glycosylated protein forms are indicated by a white dot; singly and doubly glycosylated proteins are indicated by one and two black dots, respectively. The arrowhead identifies undigested protein after PK treatment. One and two arrowheads indicate mono- and double-glycosylated protected fragments, respectively.

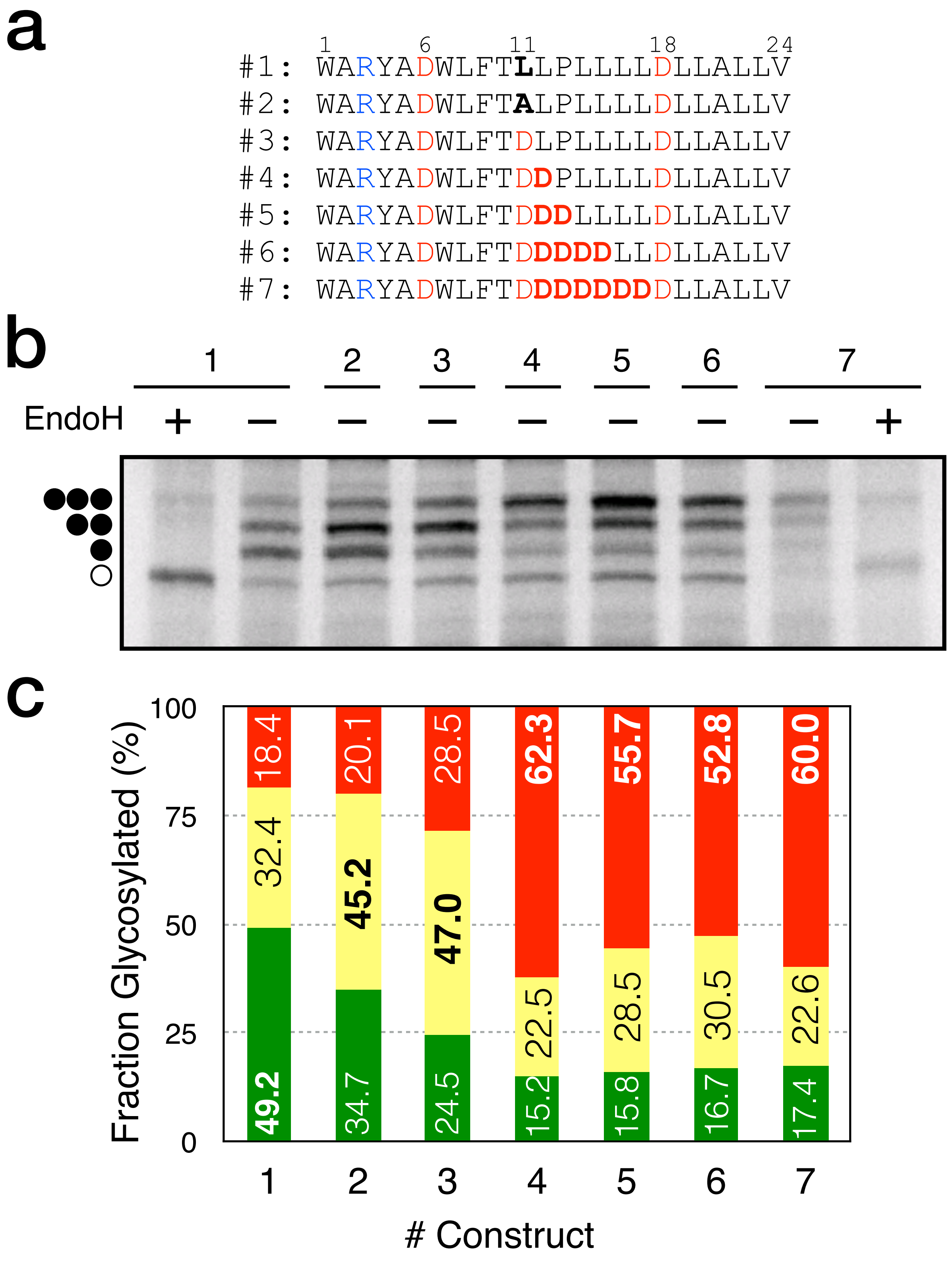


**Figure S6. Inserted/surface/translocated disposition of bRc derived sequences in the microsomal membrane. a,** Sequences of increasing hydrophilicity assayed in the LepG3 assay. Mutated residues are shown in bold. Arg and Asp residues are shown in blue and red, respectively. **b,** Representative gel of the in vitro translation of the different constructs in the presence of microsomal membranes and in the absence (—) or in the presence (+) of Endoglycosidase H (EndoH), a glycan removing enzyme. Bands of non-glycosylated protein are indicated by a white dot; single, double, and triple glycosylated proteins are indicated by one, two, or three black dots, respectively. **c,** Glycosylation profile quantification for the inserted (green), surface (yellow) and translocated (red) protein forms of each construct. The value for most abundant fraction in each construct is shown in bold.

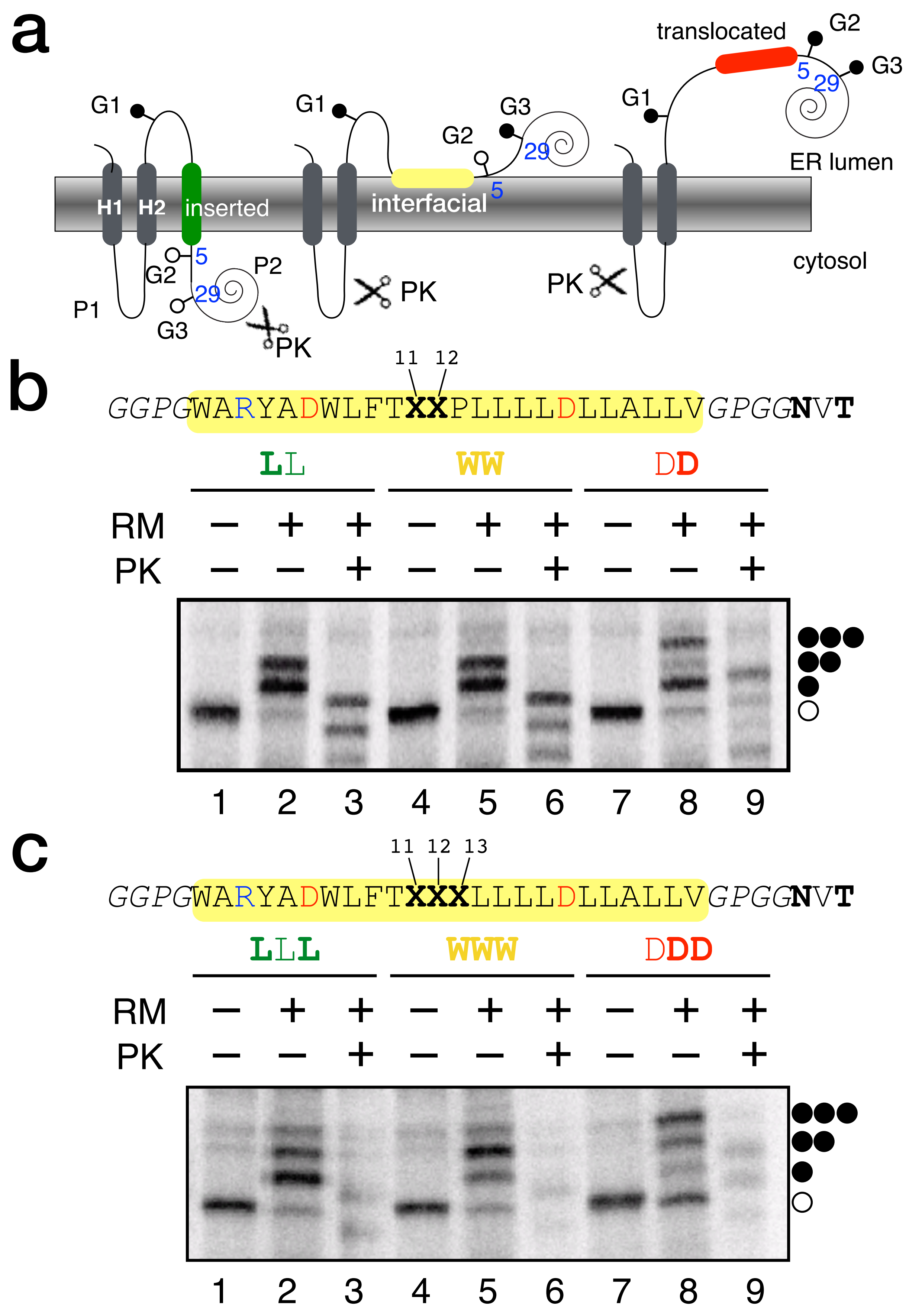


**Figure S7.** Amino acid pairs' and triplets' substitutions at central positions (Asp11 and Leu12). **a,** Schematic representation of the LepG3 design. Amino acid pairs' **(b)** and triplets' **(c)** substitutions at residues D11/L12 and D11/L12/P13. Top, background sequences used. Bottom, *in vitro* translation in the absence (—) or presence (+) of rough microsomes (RM) and/or proteinase K (PK) of bRc derived sequence harboring substituted amino acids (mutated residues in bold) at the central D11/L12 pair or D11/L12P13 triplet. Bands of non-glycosylated proteins are indicated by a white dot; single, double, and triple glycosylated proteins are indicated by one, two, or three black dots, respectively.

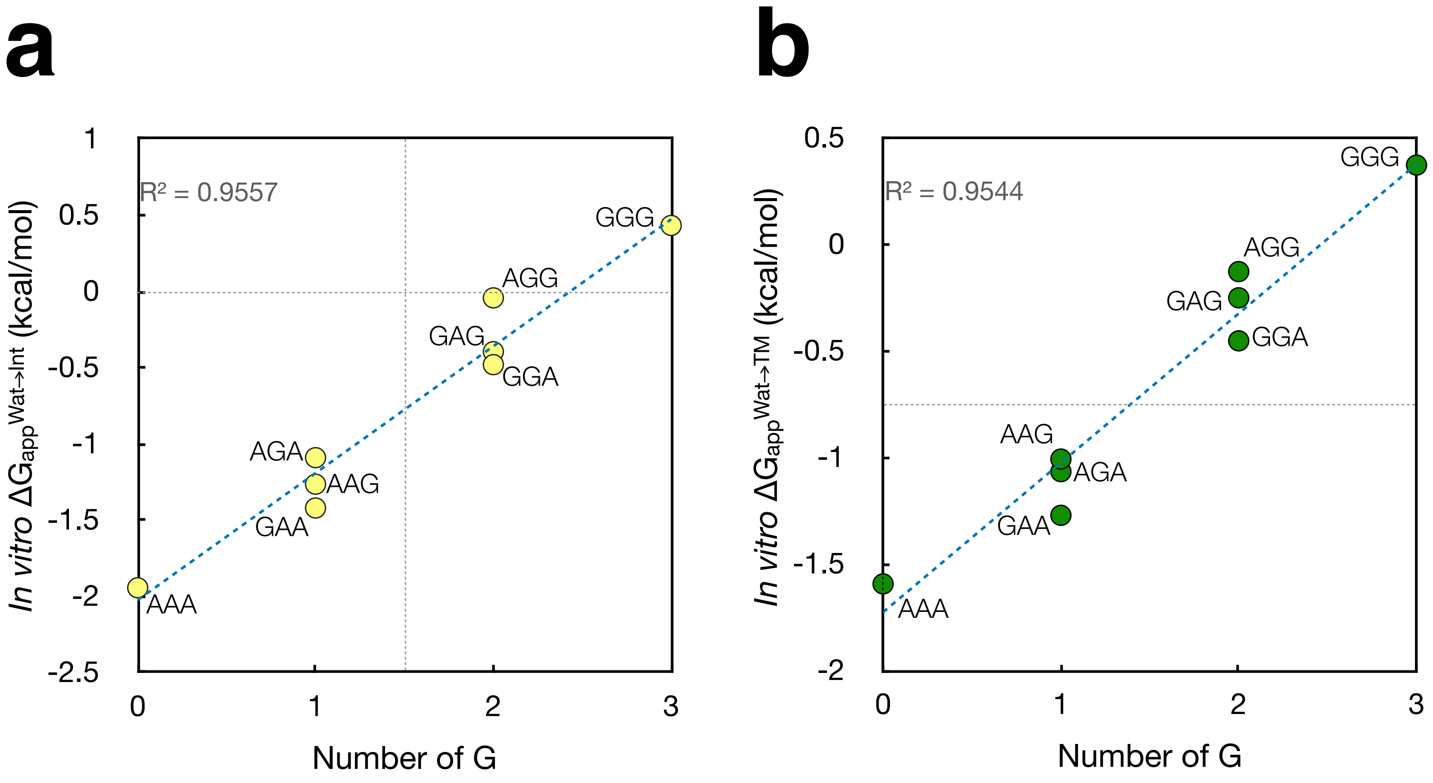


**Figure S8.** $\Delta G_{app}$ values for bRc-derived sequences containing Ala-Gly triplets. Individual points for the $\Delta G_{app}$ Water to Interface (**a**) and Water to TM (**b**) for each Ala-Gly combinations in the triplets.

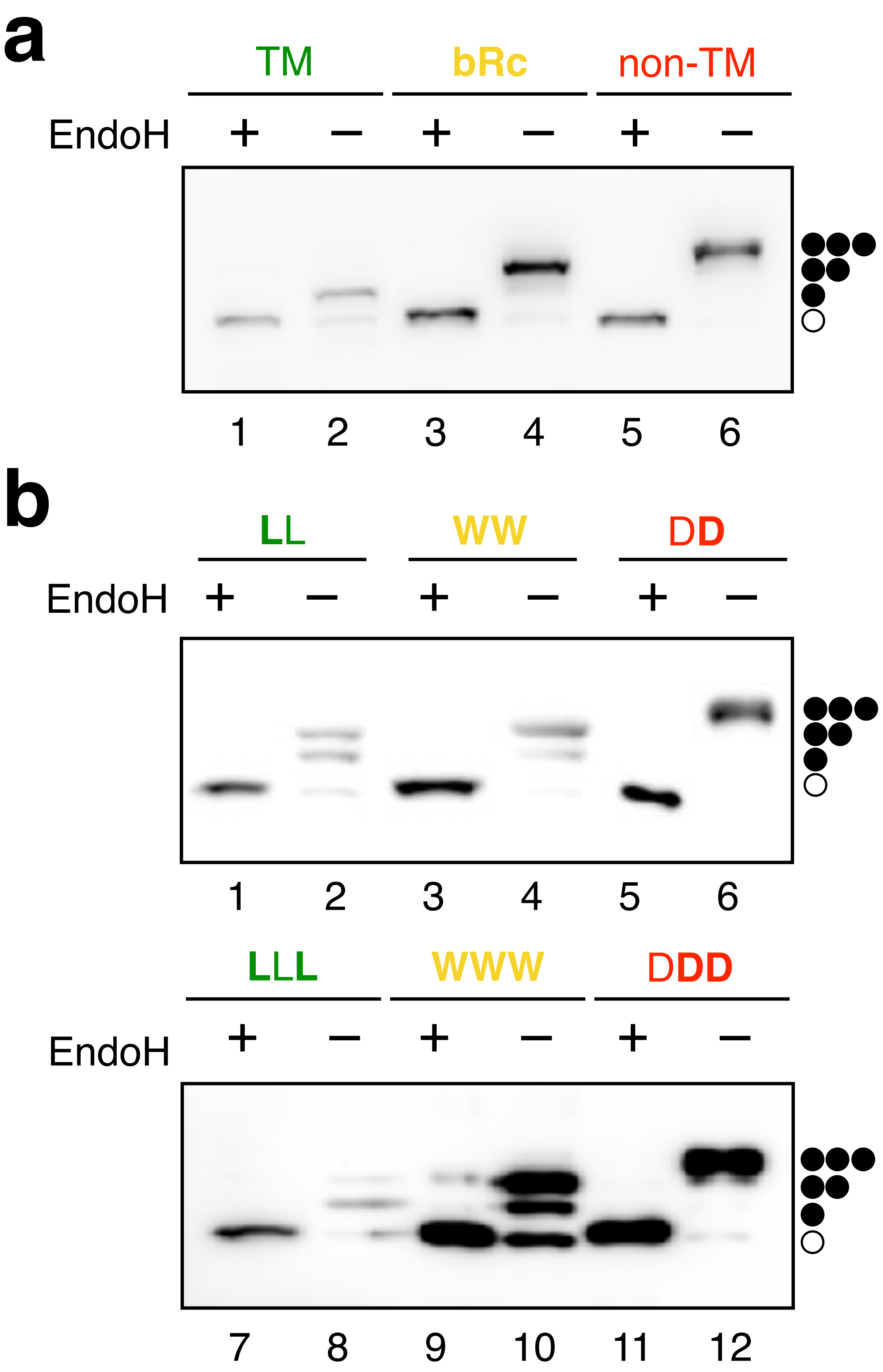


**Figure S9.** *In vivo* expression of LepG3 c-myc tagged constructs in HEK293T cells. **a,** Representative western blot of of LepG3 constructs harboring the first TM segment for *Turnip Crinkle Virus* movement protein (TM) [15], bRc derived interfacial sequence, and a translocated pseudo-randomized sequence (non-TM) from [16]. **b,** Representative western blots of the *in vivo* expression of bRc derived constructs. Top, single point mutants D11L and L12D (lanes 1-2 and 5-6, respectively) and double mutant D11W/L12W (lanes 3-4). Bottom, double mutants D11L/P13L and L12D/P13D (lanes 1-2 and 5-6, respectively) and triple mutant D11W/L12W/P13W (lanes 3-4). Before loading the samples were treated in the presence (+, lanes 1, 3, 5, 7, 9 and 11) or in the absence (—) of Endoglycosidase H (EndoH), a glycan removing enzyme. Bands of non-glycosylated protein are indicated by a white dot; singly-, doubly- and triply-glycosylated proteins are indicated by one, two and three black dots, respectively. Mutated residues are shown in bold.

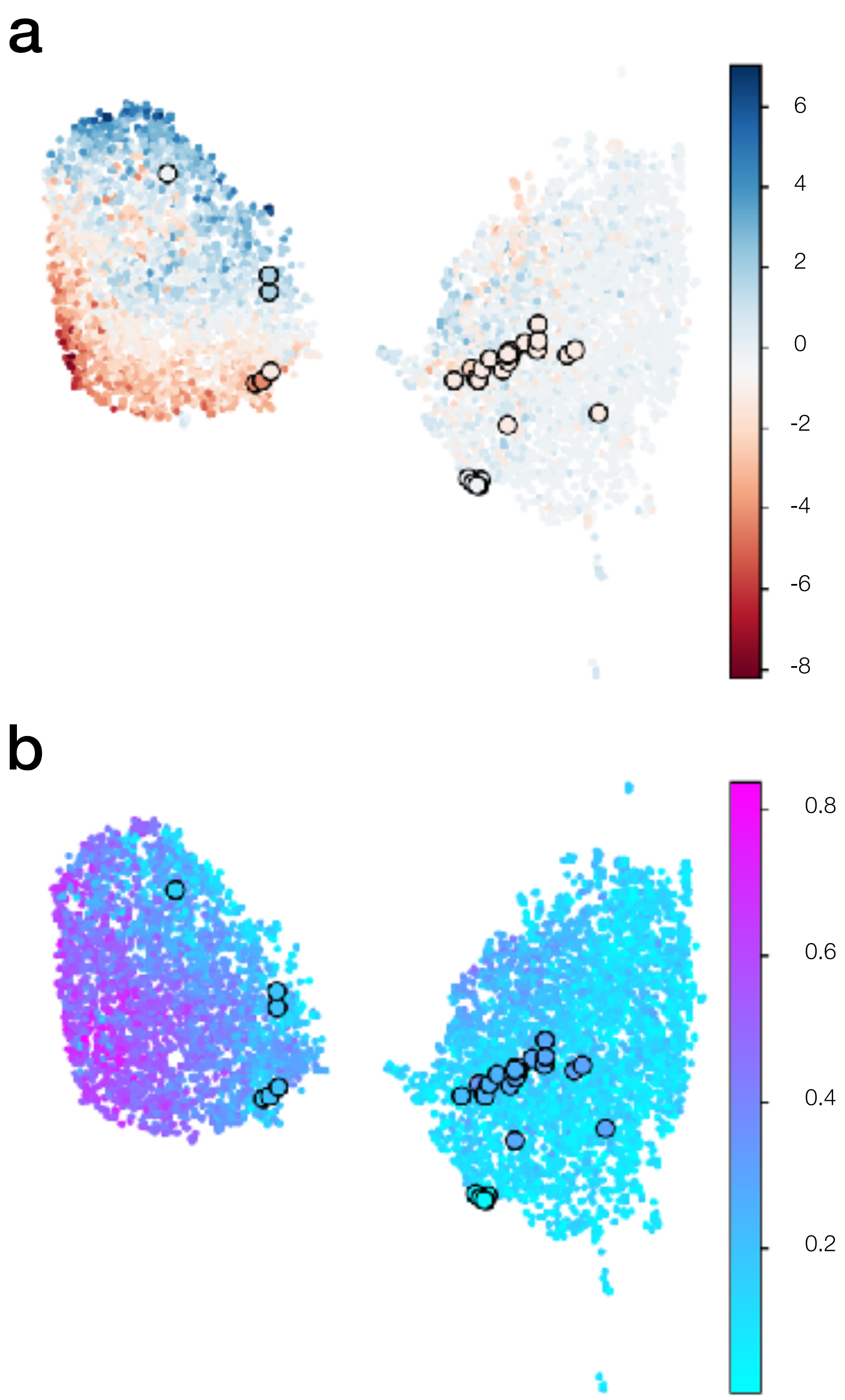


**Figure S10.** UMAP embedings as in Fig. 5. The experimentally tested sequences are embedded in the same space as large points with black outlines. **a.** colored according to net charge at pH 7, the lobe consisting mainly of helices from soluble proteins (left) shows a clear gradient from top to bottom of net charge, a pattern that is not replicated in the predominantly transmembrane lobe (right). **b.** Coloring according to hydrophobic moment, calculated using $\Delta G_{PDB}$ in a comparable manner to the hydrophobicity (Fig. 5b), shows low hydrophobic moments throughout the predominantly transmembrane lobe (right) and a clear vertical gradation among the helices from soluble proteins (left).

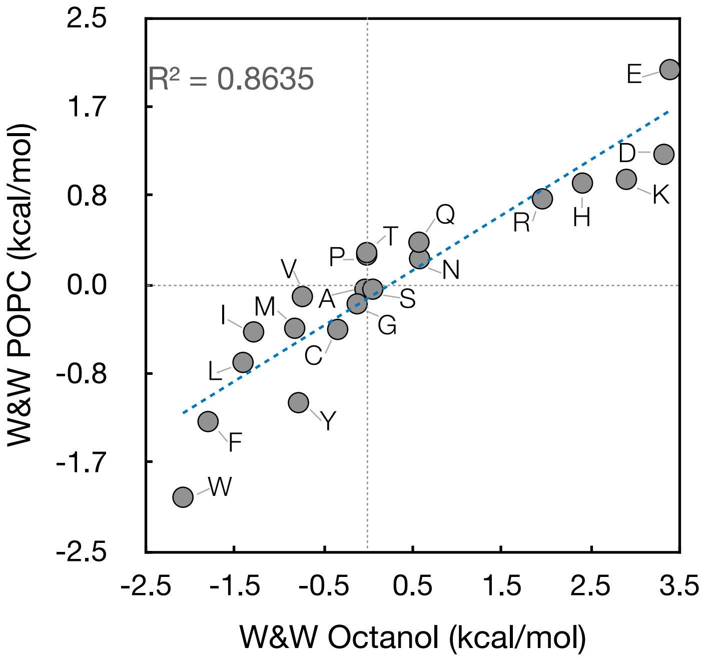


**Figure S11.** Correlation between the Wimley–White POPC (interfacial) and water/octanol (hydrophobicity) free energy scales.

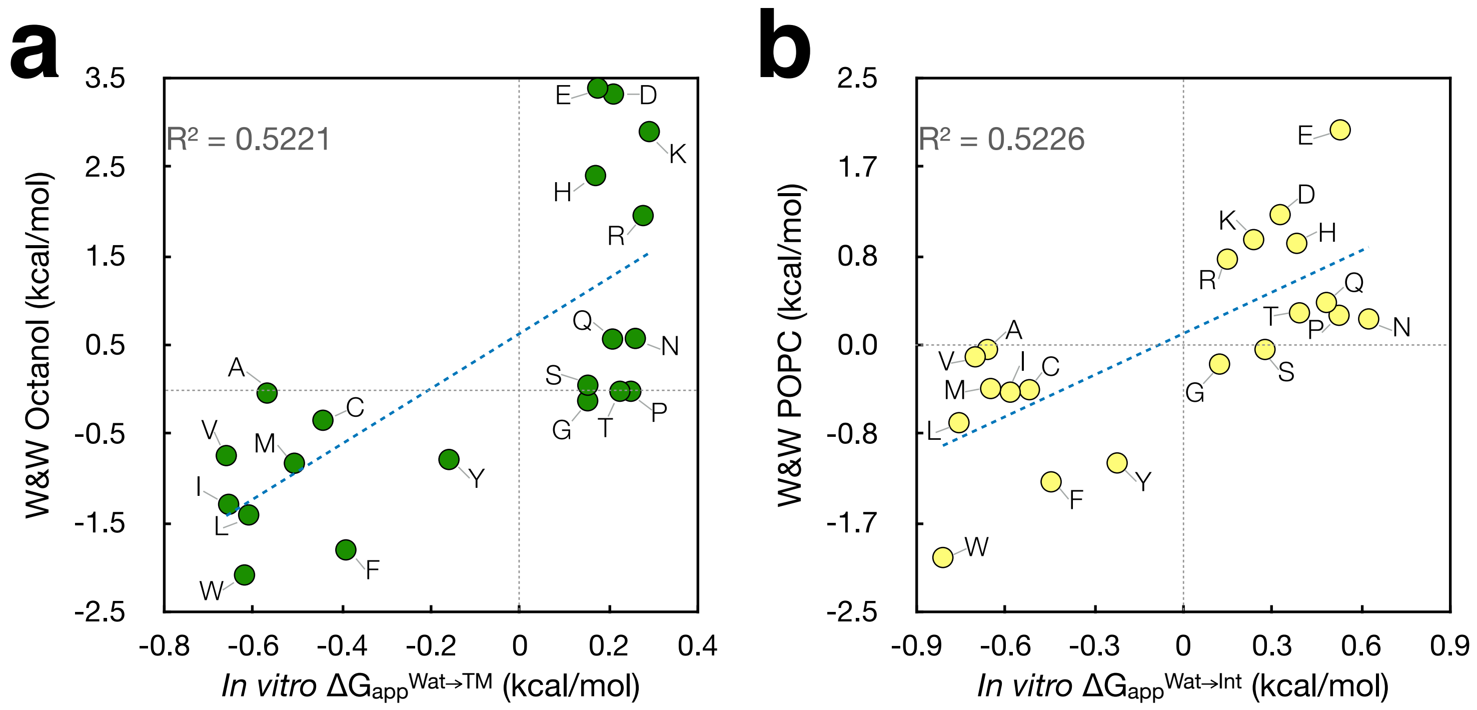


**Figure S12.** Correlations between our biological scales and the scales of Wimley and White. **a**, Correlation between the Wimley-White octane scale and our biological $\Delta G_{app}^{Wat \rightarrow TM}$ scale. **b**, Correlation between the Wimley-White POPC (interfacial) scale and our biological $\Delta G_{app}^{Wat \rightarrow Int}$ scale.

| | $\Delta G_{app}^{Wat \rightarrow Int}$ | $\Delta G_{app}^{Wat \rightarrow TM}$ |
|---|---|---|
| W | -0.81 | -0.62 |
| L | -0.76 | -0.61 |
| V | -0.70 | -0.66 |
| A | -0.66 | -0.57 |
| M | -0.65 | -0.51 |
| I | -0.58 | -0.66 |
| C | -0.52 | -0.44 |
| F | -0.45 | -0.39 |
| Y | -0.22 | -0.16 |
| G | 0.12 | 0.15 |
| R | 0.15 | 0.28 |
| K | 0.24 | 0.29 |
| S | 0.28 | 0.15 |
| D | 0.33 | 0.21 |
| H | 0.38 | 0.17 |
| T | 0.39 | 0.22 |
| Q | 0.48 | 0.21 |
| E | 0.53 | 0.25 |
| P | 0.53 | 0.17 |
| N | 0.63 | 0.26 |

**Supplementary Table 1.** *In* vitro biological $\Delta G_{app}$ scales values. Values have been derived of the evaluation of 112 measurements of a total of 27 variants translated in the *in vitro* LepG3 assay. Amino acid are represent as 1 letter code. Values are rounded off to two decimal places.

LepB P2 segment: DKQEGEWPTGLRLSRIGGI

WARYADWLFT**KKK**LLLLDLLALLV
WARYADWLFT**RRR**LLLLDLLALLV

WARYADWLFT**DDD**LLLLDLLALLV
WARYADWLFT**EEE**LLLLDLLALLV
WARYADWLFT**QQQ**LLLLDLLALLV

WARYADWLFT**DLP**LLLLDLLALLV
WARYADWLFT**HHH**LLLLDLLALLV
WARYADWLFT**NNN**LLLLDLLALLV
WARYADWLFT**PPP**LLLLDLLALLV
WARYADWLFT**SSS**LLLLDLLALLV
WARYADWLFT**TTT**LLLLDLLALLV
WARYADWLFT**YYY**LLLLDLLALLV

WARYADWLFT**GGG**LLLLDLLALLV
WARYADWLFT**AGG**LLLLDLLALLV
WARYADWLFT**GGA**LLLLDLLALLV
WARYADWLFT**GAG**LLLLDLLALLV
WARYADWLFT**AAG**LLLLDLLALLV
WARYADWLFT**GAA**LLLLDLLALLV
WARYADWLFT**AGA**LLLLDLLALLV
WARYADWLFT**AAA**LLLLDLLALLV

WARYADWLFT**CCC**LLLLDLLALLV
WARYADWLFT**MMM**LLLLDLLALLV
WARYADWLFT**VVV**LLLLDLLALLV
WARYADWLFT**WWW**LLLLDLLALLV

WARYADWLFT**FFF**LLLLDLLALLV
WARYADWLFT**III**LLLLDLLALLV
WARYADWLFT**LLL**LLLLDLLALLV

AAAAAAA**L**AAA**L**AAAAAAA
AAAAAAA**L**A**L**A**L**AAAAAAA
AAA**L**AA**L**AAAAA**L**AA**L**AAA
AAAA**L**A**L**AAAAA**L**A**L**AAAA
AAA**L**AA**L**AA**L**AA**L**AA**L**AAA

**Supplementary Table 2.** The eight clusters into which the experimentally examined sequences were split for visual examination in Figure 5a, colored using the same color scheme. Triple amino acid substitutions in the bRc-derived peptide are denoted in bold.

| | $\Delta G_{app}^{Wat \rightarrow TM}$ (kcal/mol) | Mean ± SD (kcal/mol) | $\Delta G_{app}^{Wat \rightarrow Int}$ (kcal/mol) | Mean ± SD (kcal/mol) |
|---|---|---|---|---|
| AAG | -1.101 | -1.007 ± 0.214 | -1.179 | -1.264 ± 0.119 |
| | -0.761 | | -1.250 | |
| | -0.912 | | -1.189 | |
| | -1.251 | | -1.435 | |
| AGA | -0.965 | -1.066 ± 0.227 | -1.035 | -1090 ± 0.172 |
| | -1.161 | | -1.213 | |
| | -0.744 | | -0.871 | |
| | -1.393 | | -1.241 | |
| GAA | -1.053 | -1.271 ± 0.339 | -1.200 | -1.420 ± 0.242 |
| | -1.169 | | -1.287 | |
| | -1.087 | | -1.440 | |
| | -1.1775 | | -1.752 | |
| AGG | -0.147 | -0.127 ± 0.154 | 0.269 | -0.042 ± 0.220 |
| | 0.065 | | -0.096 | |
| | -0.116 | | -0.089 | |
| | -0.311 | | -0.250 | |
| GAG | -0.334 | -0.250 ± 0.216 | -0.234 | -0.395 ± 0.153 |
| | -0.162 | | -0.477 | |
| | -0.002 | | -0.304 | |
| | -0.503 | | -0.566 | |
| GGA | -0.816 | -0.452 ± 0.301 | -0.642 | -0.480 ± 0.143 |
| | -0.311 | | -0.485 | |
| | -0.123 | | -0.293 | |
| | -0.558 | | -0.500 | |

**Supplementary Table 3.** Apparent partitioning free energies for replicates of the sequences containing both Ala and Gly, alongside the mean and standard deviations of the free energies across the replicates.
